# Phenotypic and functional characterization of corneal endothelial cells during in vitro expansion

**DOI:** 10.1101/717405

**Authors:** Ricardo F. Frausto, Vinay S. Swamy, Gary S. L. Peh, Payton M. Boere, E. Maryam Hanser, Doug. D. Chung, Benjamin L. George, Marco Morselli, Liyo Kao, Rustam Azimov, Jessica Wu, Matteo Pellegrini, Ira Kurtz, Jodhbir S. Mehta, Anthony J. Aldave

## Abstract

The advent of cell culture-based methods for the establishment and expansion of human corneal endothelial cells (CEnC) has provided a source of transplantable corneal endothelium, with a significant potential to challenge the one donor-one recipient paradigm. However, concerns over cell identity remain, and a comprehensive characterization of the cultured CEnC across serial passages has not been performed. To this end, we compared two established CEnC culture methods by assessing the transcriptomic changes that occur during in vitro expansion. In confluent monolayers, low mitogenic culture conditions preserved corneal endothelial cell state identity better than culture in high mitogenic conditions. Expansion by continuous passaging induced replicative cell senescence. Transcriptomic analysis of the senescent phenotype identified a cell senescence signature distinct for CEnC. We identified activation of both classic and new cell signaling pathways that may be targeted to prevent senescence, a significant barrier to realizing the potential clinical utility of in vitro expansion.

## INTRODUCTION

The worldwide shortage of donor corneal tissue for the treatment of corneal endothelial dysfunction necessitates the development of viable alternatives to the paradigm of one donor cornea being used for only one recipient (Mehta et al., 2019; Okumura et al., 2014b; Soh et al., 2017). Corneal endothelial cell (CEnC) dysfunction is the primary indication for corneal transplantation both in the United States and worldwide, and while endothelial keratoplasty represents a significant advance in the surgical management of corneal endothelial dysfunction, the worldwide shortage of surgical-grade donor corneas and the lack of adequately trained surgeons in the majority of countries, as well as a variety of associated intraoperative and postoperative complications, have significantly limited the impact of endothelial keratoplasty on visual impairment worldwide due to corneal endothelial dysfunction (Deng et al., 2015; Lass et al., 2017; Van den Bogerd et al., 2018). The in vitro generation of stem cell-derived corneal endothelial-like cells (Yamashita et al., 2018), immortalized CEnC lines (Schmedt et al., 2012; Valtink et al., 2008) and expansion of primary CEnC from cadaveric donor corneal tissue (Okumura et al., 2014b; Parekh et al., 2016; Peh et al., 2019) have challenged the one donor-one recipient paradigm of corneal transplantation. Nevertheless, in vitro culture poses its own challenges, including unwanted changes in cell phenotype (e.g., endothelial to fibroblastic) and progression towards replicative senescence that limits cell numbers (Sheerin et al., 2012; Soh et al., 2017). In addition, the quality of the donor tissue from which the CEnC are derived is critical in the successful establishment of an in vitro CEnC culture. Donor age significantly impacts culture success rate, with the optimal age being less than 40 years old. Reduced success rates from older donors are correlated with an appearance of senescence-associated markers ex vivo (Mimura and Joyce, 2006), and this is believed to significantly limit re-entry into the cell cycle even in the presence of potent mitogenic factors. While senescence is generally an irreversible cell state (Campisi, 2013), CEnC from younger donors are characterized by a quiescent cell state (G1 cell cycle arrest), which is a reversible mitotic arrest that enables these cells to undergo mitogen-induced cell cycle re-entry (Joyce, 2005). Donor age and other factors (e.g., days in preservation medium, donor medical history, cell count) dictate the success of establishing in vitro cultures (Soh et al., 2017). This makes identifying an optimal in vitro culture protocol essential for ensuring consistent establishment and expansion of CEnC.

The corneal endothelium is a neuroectoderm-derived tissue that is located on the posterior surface of the cornea and is a semipermeable monolayer of mitotically inactive (i.e., quiescent) CEnC. A critical functional property of the corneal endothelium is to maintain the corneal stroma in a relatively dehydrated state. This process ensures that the collagen fibers of the stroma retain an ultrastructural organization essential for corneal transparency. The pump-leak hypothesis is believed to best explain the role that the endothelium plays in maintaining a relatively dehydrated stroma. Passive movement of water from the aqueous humor to the stroma (i.e., leak) and active transport of solutes (i.e., pump) in the opposite direction are regulated by the CEnC barrier formation (i.e., tight junctions, cell adhesion) and expression of solute transporters (e.g., Na/K ATPases, SLC4A11) (Gottsch et al., 2003). Mutations in genes associated with transporter function (e.g., *SLC4A11*) or CEnC identity (e.g., *ZEB1, TCF4*) lead to stromal edema and loss of corneal clarity (Baratz et al., 2010; Krafchak et al., 2005; Vithana et al., 2006; Wieben et al., 2012). In general, loss of barrier integrity, dysfunction of solute transporters or a significant decrease in CEnC density leads to loss of corneal clarity that necessitates corneal transplantation. Endothelial cell failure constitutes the primary indication for corneal transplantation in the U.S., serving at the indication for 55% of all keratoplasty procedures performed in 2018 (2018). While endothelial keratoplasty remains the primary method for managing endothelial cell dysfunction in the U.S., the aforementioned factors that have limited the impact of endothelial keratoplasty on decreasing the global burden of vision loss from endothelial dysfunction necessitates the development of alternative therapeutic interventions.

To achieve this goal, we assessed two previously reported methods for establishing cultures of primary CEnC (Bednarz et al., 1998; Peh et al., 2015) by using a multipronged approach. We determined the impact of in vitro expansion on CEnC gene expression by performing a transcriptomics analysis, and identified gene expression features of replicative senescence. In addition, we performed a variety of assays to determine the impact of these two methods on essential CEnC functions. We identified new potential targets for suppressing cellular senescence, and confirmed that a relatively low mitogenic environment is better at maintaining the CEnC phenotype in vitro (Peh et al., 2015). These findings form the basis for continued development of in vitro culture and expansion of primary CEnC for their eventual use in cell replacement therapy for the management of corneal endothelial loss or dysfunction.

## METHODS

### Primary corneal endothelial cell cultures

Corneas used in this study were obtained from commercial eye banks (Table S1). Criteria used for selection of high quality donor corneal tissue were: 1) donor younger than 40 years (mean: 17.6; range: 2- 35); 2) no donor history of diabetes or corneal disease; 3) endothelial cell density greater than 2300 cells/mm^2^ (mean: 3019; range: 2387-3436); 4) death to preservation less than 12 hours, if body not cooled, or less than 24 hours, if body cooled; and 4) death to culture less than 15 days (mean: 6; range: 2- 14). Descemet membrane with attached endothelium was stripped from the stroma using the method commonly employed in preparation of the donor cornea for Descemet membrane endothelial keratoplasty. Seven independent CEnC cultures were established using two previously described protocols with minor modifications (Bednarz et al., 1998; Peh et al., 2015). One method utilizes trypsin for dissociation of endothelial cells from Descemet membrane, followed by seeding on laminin coated cell culture plastic, and culturing in a 1:1 mixture of F12-Ham’s and M199 (F99) medium. The second method utilizes collagenase A for dissociation of endothelial cells from Descemet membrane. This is followed by seeding on collagen IV coated cell culture plastic, and culturing initially in Endothelial SFM (M5) followed by culturing in a 1:1 mixture of F12-Ham’s and M199 (M4) medium. When cells reach confluence, the medium is changed back to M5 medium for establishment and maintenance of the CEnC phenotype. Cell passaging is performed with TrypLE Select (Thermo Fisher Scientific). Cells isolated using each protocol are referred to as F99 cells or M5 (M4/M5 or dual media) cells to indicate the method used to establish the cultures.

### Total RNA isolation

Primary CEnC were lysed in TRI Reagent (Thermo Fisher) and total RNA was prepared as per the manufacturer’s instructions. RNA preparations were subsequently purified using the RNeasy Clean-Up Kit (Qiagen, Valencia, CA). The quality of the total RNA was assessed with both the Agilent 2100 Electrophoresis Bioanalyzer System (Agilent Technologies, Inc., Santa Clara, CA) and the Agilent TapeStation 2200 (Agilent Technologies, Inc.).

### RNA-sequencing and data processing

RNA was isolated and RNA-seq libraries were prepared using the KAPA mRNA HyperPrep Kit with an automated liquid handler (Janus G3 – PerkinElmer) according to the manufacturer’s instructions. Library preparation was performed at the UCLA Institute for Quantitative and Computational Biology. DNA libraries were submitted to the UCLA Technology Center for Genomics and Bioinformatics for sequencing, which was performed on the Illumina HiSeq 3000 platform. All RNA-seq data were single-end 50 base reads. Reads were aligned to the human GRCh38.p12 genome, and transcripts were quantified using the kallisto (v0.44.0) program (Bray et al., 2016) with the Ensembl Annotation Release version 92. Quantities were given in transcripts per million (TPM), and differential gene expression analysis was performed with the Sleuth (v0.30.0) R-package (Pimentel et al., 2017). Differential expression was tested using a likelihood ratio test (negative binomial test), and corrected for multiple testing using the Benjamini-Hochberg correction. Given the sporadic availability of donor corneas, each of the seven cultures was established as an individual batch. To account for batch effects in the data, we included batch number (i.e., culture number) as a covariate in the model used to test for differential expression. The following thresholds defined differential expression: fold change>1.5, TPM>11.25 and q-value<0.05. Of note, our background threshold was 7.5 TPM, and thus the TPM threshold of 11.25 represents a 1.5 fold change increase above the background threshold. With the goal of achieving a balance between Type I and Type II errors, the background threshold was selected to retain genes with low expression (e.g., *ZEB1* with a TPM of ∼25), but was sufficiently robust to exclude many genes with low (<7.5 TPM) abundance values. RNA-seq data generated from ex vivo corneal cell types were obtained from the GEO DataSets database (accession GSE121922). RNA-seq data generated for this study were submitted to GEO DataSets and assigned accession number GSE132204.

### Gene ontology and pathway analysis

Gene ontology (GO) and pathway analysis on gene symbols only was performed using the web-based tool gProfiler (version: e94_eg41_p11_04285a3). The three main GO categories are Molecular Function (MF), BcCellular Component (CC) (data version: release/2018-12-28). Databases used for pathway analysis were Kyoto Encyclopedia of Genes and Genomes (KEGG) pathways (version: KEGG FTP Release 2019-01-07), REACTOME (REAC) pathways (classes: 2019-1- 24), and WikiPathways (WP)(version: 20190110). The g:SCS method for computing multiple testing correction was used for selecting significantly enriched GO terms and pathways identified by gProfiler. Ingenuity Pathway Analysis software (Qiagen; build: 486617M) was used to determine the pathways enriched and activated in our lists of differentially expressed genes.

### Quantitative PCR

Quantitative polymerase chain reaction (qPCR) was performed to validate gene transcript abundances observed by RNA-seq. Total RNA (100 ng) was subjected to first-strand cDNA synthesis using the SuperScript III First-Strand Synthesis kit (Thermo Fisher Scientific) with oligo-dT primers. Reactions were performed on the LightCycler 480 System (Roche) using the KAPA SYBR FAST qPCR Kit (Roche) and gene-specific oligonucleotide primers obtained from the Harvard Primer Bank database (Spandidos et al., 2008; Spandidos et al., 2010; Wang and Seed, 2003) or designed using the NCBI Primer-BLAST tool (Table S2). Relative transcript abundances were calculated in comparison with the housekeeping gene *RAB7* using the comparative Ct (2^-ΔCt^) method (Livak and Schmittgen, 2001). Transcript quantities were graphed as 2^-ΔCt^.

### Capillary-based, automated protein immunodetection assay

Cells were lysed with RIPA buffer and whole-cell lysates were prepared and processed for protein detection using the Wes separation 12-230 kDa capillary cartridges (Protein Simple). Separation and detection were performed as per the manufacturer’s instructions. Quantification and data analysis were performed using the Compass for SW software (version 3.1.7; build ID: 1205). Primary antibodies are listed in Table S3).

### Morphometric analysis

At each passage, we acquired phase contrast images of confluent monolayers established in either F99 or M5. Image acquisition was performed with the Leica DMIL LED inverted microscope (Leica Microsystems), the N PLAN L 20x/0.35 PH1 objective, and the SPOT Insight color camera (Diagnostic Instruments, Inc.). Images were captured with the SPOT software (version 4.6). Image analysis was performed using ImageJ 1.51h (National Institutes of Health). We created regions of interest (ROI) by outlining the periphery of at least 20 representative cells using the freehand tool. ImageJ output the values for area (um2) and circularity (ratio) for the selected ROI.

### Cell barrier assay

The electrode array 8W10E+ ECIS (Applied BioPhysics) was stabilized with F99 or M5 medium. The array surface was coated with 40 µg/cm^2^ chondroitin sulfate A (Sigma-Aldrich) and 400 ng/cm^2^ laminin (Sigma-Aldrich) in phosphate-buffered saline (PBS) for two hours (for F99 cells) or with 7.5 µg/cm^2^ collagen IV (Sigma-Aldrich) in 1x HEPES for 30 min (for M5 cells). Cells were seeded at a density to achieve 100% confluence shortly after seeding, and were incubated in the arrays at room temperature for one hour to facilitate even distribution. After seeding and preliminary cell attachment, arrays were positioned into a 16-well array station and connected to the ECIS Zθ instrument to measure electric impedance (Ω at 4000 Hz) for 4 days. Cell-cell (*R*_b_, Ω • cm^2^) and cell-substrate (*α*, Ω^1/2^ • cm) adhesion were modeled from the electric impedance data obtained at 4000 Hz (Stolwijk et al., 2015).

### Transporter assays

Lactate (SLC16A1), bicarbonate (SLC4A4) and proton (SLC4A11) transport were monitored by measuring free H^+^ concentration (pHi) with a microscope fluorometer (Kao et al., 2016) using the fluorescence-based (dual-excitation 500 nm and 440 nm) ratiometric pH indicator BCECF (Thermo Fisher Scientific), which was pre-loaded into the cells prior to substrate exposure.

#### SLC16A1 transporter

BCECF loading was performed in lactate-free solution (20mM Na gluconate, 120mM NaCl, 1mM CaCl_2_, 1mM MgCl_2_, 2.5mM K_2_HPO_4_, 5mM dextrose and 5mM HEPES, pH 7.4). Fluorescence was monitored until a stable pH_i_ was achieved. Subsequently, the lactate-free buffer was replaced by perfusion with lactate-containing solution (20mM Na lactate in place of 20mM Na gluoconate) for about 200 seconds and then switched back to the lactate-free solution.

#### SLC4A4 transporter

BCEFC loading was performed in bicarbonate-free solution (140mM TMACl, 2.5mM K2HPO4, 1mM CaCl2, 1mM MgCl2, 5mM Dextrose and 5mM HEPES, pH 7.4). After pH_i_ stabilization was attained, the solution was replaced by perfusion with a solution containing tetramethylammonium bicarbonate (115mM TMACl and 25mM TMAHCO3 in place of 140mM TMACl and 5mM HEPES). Perfusion was performed for about 200 seconds and then switched to a solution containing sodium bicarbonate (115mM NaCl, 2.5mM K2HPO4, 1mM CaCl2, 1mM MgCl2, 5mM Dextrose and 25mM NaHCO3, pH 7.4), and fluorescence was monitored for an additional 400-600 seconds.

#### SLC4A11 transporter

BCECF loading was performed in HEPES buffer (110mM KCl, 35mM TMACl and 5mM HEPES, pH 7.4). Once pH_i_ stabilized, a second HEPES buffer (110 mM KCl, 35 mM TMACl and 5 mM HEPES pH 6.2) was replaced by perfusion and pH_i_ was monitored for approximately 800 seconds.

### Cell migration assay

A non-wounding method to assess cell migration was used. Two-well silicone inserts (ibidi GmbH), each creating a 500um gap, were placed onto cell culture treated plastic. Wells were coated as described above for F99 and M5 cells. After cell isolation was performed, cells were seeded into each well and allowed to grow to confluence. Cell migration was initiated by removing the insert and was monitored for up to 60 hours by phase-contrast microscopy. Images were acquired using the Leica DMIL LED inverted microscope (Leica Microsystems) and the N PLAN L 20x/0.35 PH1 objective. Image capture was performed with the Leica DFC3000 G monochrome camera controlled with the Leica Application Suite X software (version 3.0.3.16319). Gap closure was measured at 22 hours and cell size was measured at 60 hours using ImageJ 1.51h software (National Institutes of Health).

## RESULTS

### In vitro expansion of CEnC induces senescence-associated morphogenesis

The morphogenic effects of culture in high mitogenic (F99) and low mitogenic (M5) conditions on primary CEnC were examined (Fig. 1). Phase contrast images were acquired at each passage when confluent monolayers were established (Fig. 1B and Fig. S1). Morphometric analysis was performed at each passage (Fig. 1C). Up to passage 3, the area occupied by each cell was greater in F99, compared with cells in M5, but the effect of medium on the curves was not statistically significant (p=0.065). Cell circularity, which measures the degree to which a cell shape resembles a circle (1.0 is a perfect circle), was greater at all passages for cells in M5 medium, compared with cells in F99. The effect of medium on the curves for circularity was statistically significant (p=0.042). As the value approaches 0, cell shape is increasingly irregular and/or elongated.

**Figure 1.**
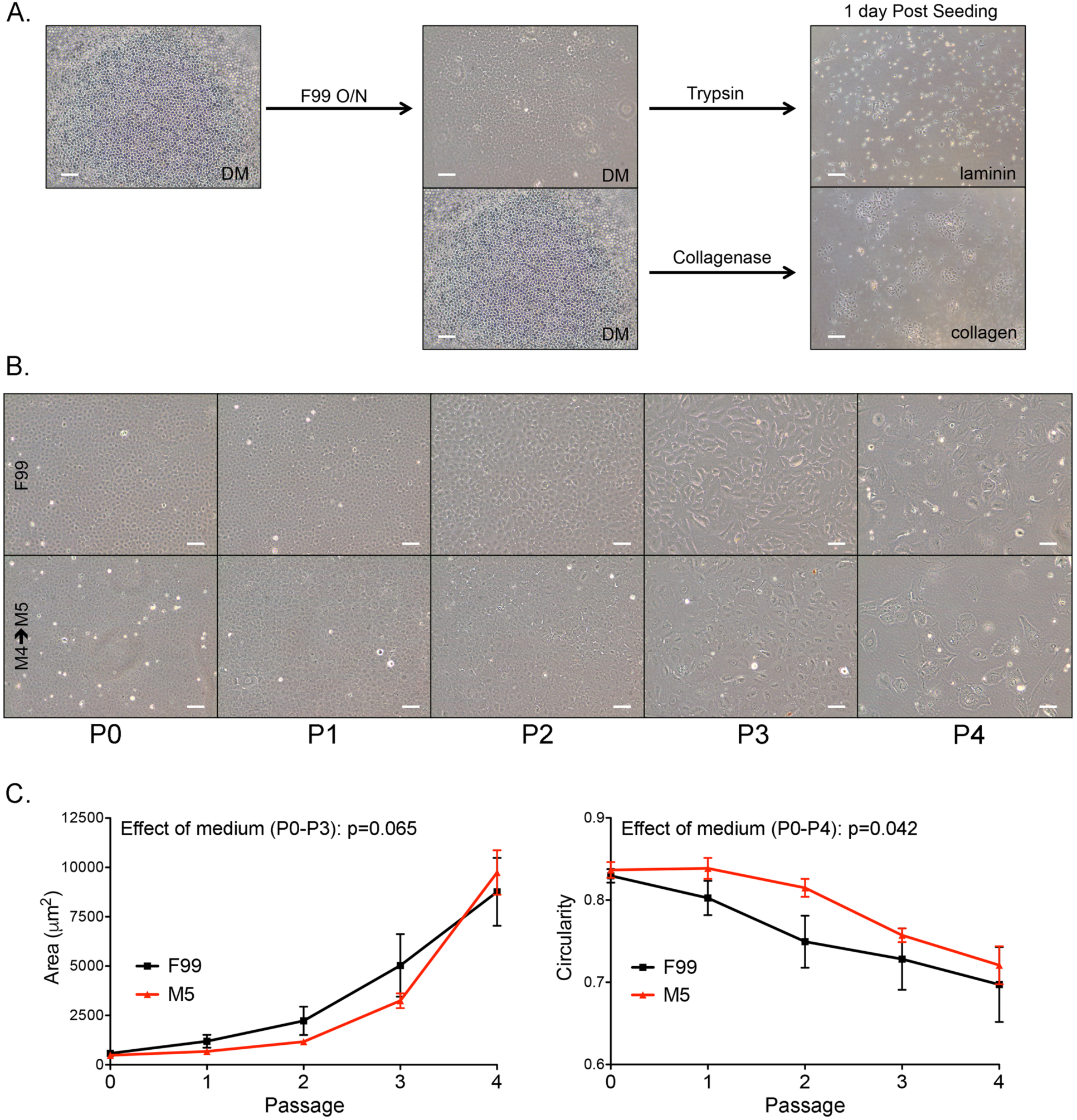
M5 medium delays morphologic features associated with a senescent phenotype. (A) Two protocols for the isolation and culture of primary CEnC. After detachment from the cornea, Descemet membrane (DM) with the attached endothelium was either incubated overnight in F99 medium at 37C (left panel) or subjected directly to collagenase digestion (middle panel). Cells incubated overnight were detached from DM with trypsin and seeded onto laminin-coated plastic. Cells dissociated from DM using collagenase were seeded onto collagen-coated plastic. Images show cells 1-day after seeding (right panel). (B) Images show confluent CEnC cultures at five passages using two culture methods (F99 or M5). (C) Line graph shows mean cell area (μm) at each passage. (D) Line graph shows mean circularity at each passage. Data in (C) and (D) are represented as the mean ±SEM (n=6). Statistical comparisons were performed using two-way ANOVA, with passage and medium defining the variables for this comparison. Scale bars, 100 μm.

### A low mitogenic environment maintains a robust CEnC-specific gene expression profile in primary CEnC

To examine the ability of the cultured cells to maintain a CEnC-specific gene expression profile in low- or high-mitogenic environments, we compared the expression of 97 genes, previously identified as specific to ex vivo corneal endothelium (evCEnC), in primary CEnC in M5 versus F99 at each passage (Fig. 2) (Frausto et al., 2016). At P0, the cells in M5 expressed 83 of the 97 (85.6%) evCEnC-specific genes, while cells in F99 expressed 76 (78.4%) (Fig. 2A). By P4, the percentages decreased to 75.3% (73/97) in cells cultured in M5, and 66% (64/97) in cells cultured in F99 (Fig. 2A). We examined the 97 evCEnC-specific genes in the passaged cells to identify those that may be suitable positive selection markers for assessing the quality of CEnC cultures using the following criteria: 1) expression in 2 or more sequential passages, starting with P0; 2) decreasing expression over increasing passages (Fig. 2B); and 3) encoding a cell surface protein. Only *TMEM178A* demonstrated these characteristics (Fig. 2C), and its expression was validated by qPCR at P0 and P3 (Fig. 2D).

**Figure 2.**
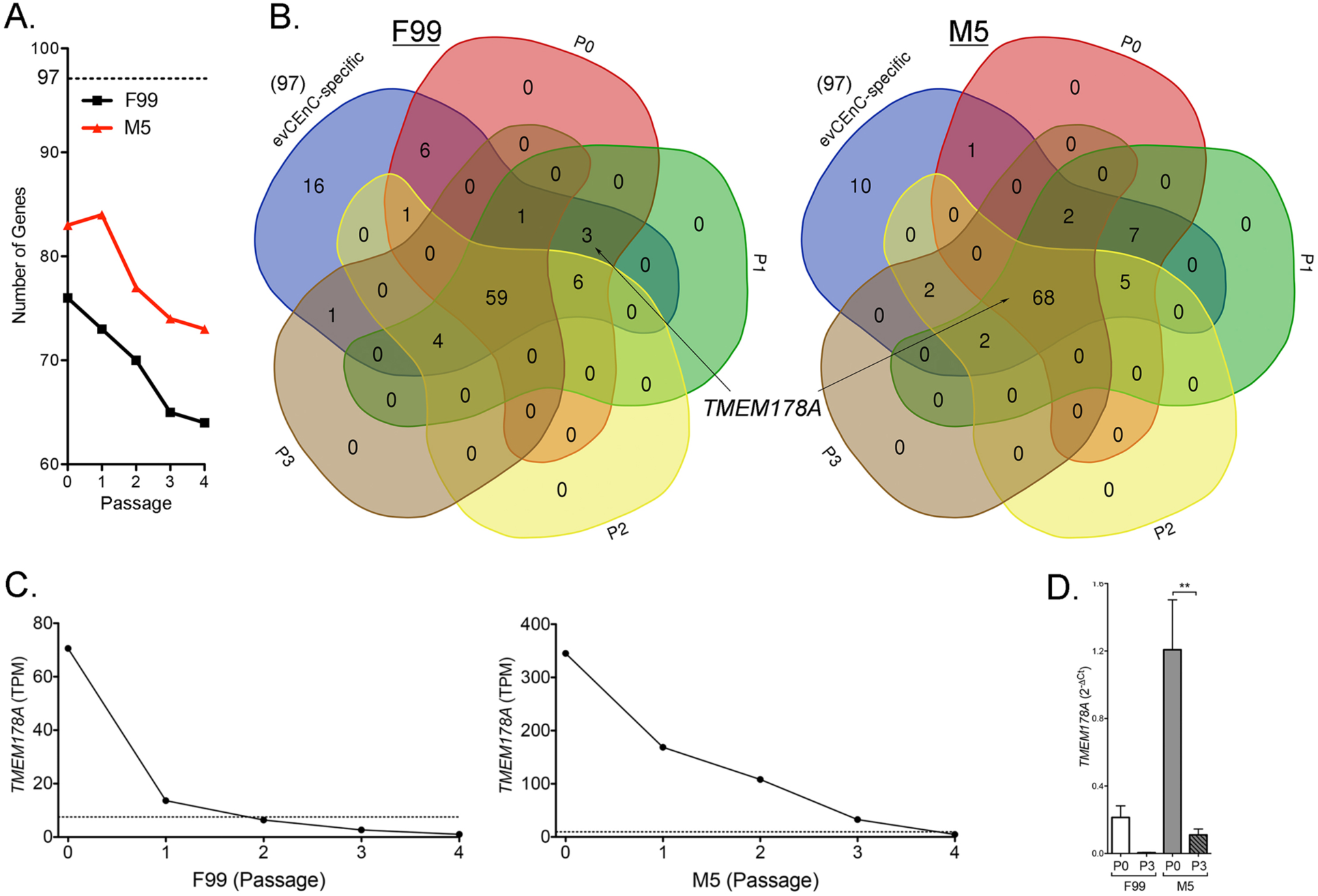
M5 medium maintains CEnC-specific gene expression in cultured CEnC. (A) Line graph shows the number of evCEnC-specific genes expressed in CEnC cultured in either F99 or M5 medium. Dashed line represents the number (97) of evCEnC-specific genes identified in a previous study. (B) Venn diagrams demonstrating the expression of evCEnC-specific genes at passages P0-P3 (P4 not shown due to difficulty visualizing Venn diagrams of greater than 5 sets). *TMEM178A* was identified as a cell surface marker expressed in early passages (at least P0 and P1) in CEnC cultured in either F99 or M5. (C) Line graphs show expression pattern of *TMEM178A* in CEnC cultured in F99 or M5. Gene was considered expressed if TPM value was greater than 7.5 (dashed line). (D) Bar graph shows *TMEM178A* transcript abundances in P0 and P3 CEnC using qPCR. Data in (D) are represented as the mean ±SEM (n=4). Statistical comparisons were performed using one-way ANOVA with post-hoc Tukey test. **, P<0.01.

An analysis of the expression of genes purported to be markers for CEnC identity and/or to determine the quality of CEnC in culture revealed that many were neither specific nor highly expressed in corneal endothelium (e.g., *ATP1A1*, *TJP1*, *VIM*, *PRDX6*, *SLC3A2*), or did not correlate well with other quality metrics, such as cell morphology (e.g., *ALCAM1*, *ERBB2*, *CD248*, *SLC25A11*)(Fig. S2). We should note that our data represents transcript abundance, and these markers may prove to be adequate based on protein abundance, which was not determined for the majority of the reported markers in this study. Based on transcript abundance, *SLC4A11* and *CD44* may represent the optimal markers for selection of high quality cultured CEnC, with *SLC4A11* classified as a positive selection marker and *CD44* classified as a negative selection marker (Fig. 3). *SLC4A11* and *CD44* transcript abundances at P0 and P3 in CEnC cultured in both F99 and M5 was determined by RNA-seq and validated by both qPCR and a Western assay (Fig. 3A). Expression of *SLC4A11*, which was high in evCEnC, decreased with increasing passage, while expression of *CD44*, which was high in evCEpC, increased with increasing passage (Fig. 3B).

**Figure 3.**
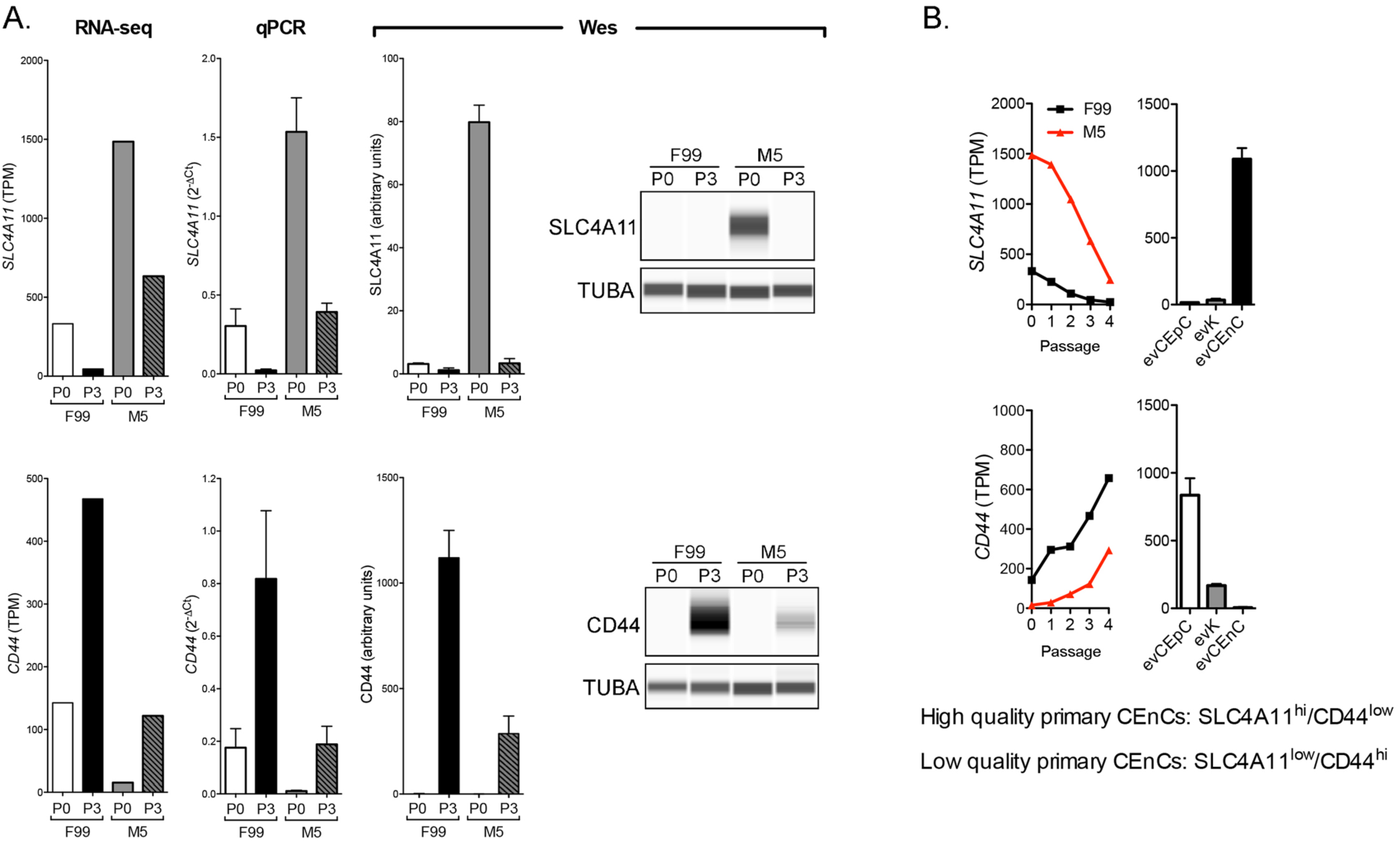
SLC4A11 and CD44 as cell surface markers for the selection of high quality cultured CEnC. Assessment of previously published markers and results obtained in this study provide evidence that SLC4A11 and CD44 represent the most informative markers for the selection of high quality cultured cells, and represent a minimum for selection. (A) Bar graphs show transcript (RNA-seq and qPCR) abundance of *SLC4A11* and *CD44* in P0 and P3 CEnC cultured in F99 and M5. Detection and quantification of SLC4A11 and CD44 was performed using a Western assay (Wes). Bar graphs show protein quantification. (B) Line graphs show *SLC4A11 and CD44* transcript quantities in CEnC at each passage (P0 – P4). Bar graphs show transcript abundance in the three main cell types of the cornea ((epithelial cells (evCEnC), keratocytes (evK) and endothelial cells (evCEnC)). Data in bar graphs in (A) are represented as the mean ±SEM (n=4, mRNA; n=3, protein). Data in line graphs are represented as the mean TPM at each passage (n=7). Data in bar graphs in (B) are represented as the mean TPM,±SEM (n=3).

### CEnC grown in a low-mitogenic environment possess a robust cellular respiration phenotype

We performed bioinformatics analyses using differentially expressed genes to determine the biological characteristics of cells cultured with F99 and M5. We identified the genes that were significantly differentially expressed in cells at each passage between the two growth conditions (Table S4). With progressive passaging, the number of genes showing differential expression (M5 compared to F99 at each passage) decreased for cells in each media condition, with the transcriptomes of the cells grown in F99 and cells grown in M5 becoming increasingly more similar (Fig. S3A). As the gene expression difference was greatest at P0 (between F99 and M5), we performed bioinformatics analyses on the data set obtained at P0. The genes upregulated in cells with each media at P0 were subsequently analyzed to identify gene ontology (GO) and pathway terms enriched in our data sets (Table S5). GO terms significantly (q-value <0.05) enriched in the F99 (P0) upregulated genes data set were associated with cell differentiation and tissue development (e.g., developmental process, system development, and multicellular organism development)(Table 1). In addition, significantly enriched pathway terms were associated with cell state transitions (e.g., TGF-beta signaling pathway and epithelial to mesenchymal transition in colorectal cancer) and cell senescence (e.g., senescence and autophagy in cancer). The significantly enriched GO terms in the M5 (P0) upregulated genes data set were associated with cellular respiration (e.g., oxidoreductase activity and electron transfer activity) and lipid metabolism (e.g., lipid oxidation and fatty acid oxidation). Similarly, pathway terms enriched in this data set were also associated with cellular respiration (e.g., electron transport chain (OXPHOS system in mitochondria)) and lipid metabolism (e.g., fatty acid metabolism). GO and pathway enrichment results were similar for the other passages within the same media group (Table S5).

**Table 1.** Gene ontology and pathway enrichment analysis of the P0 inter-media comparison.

| Source | Term Name | Term ID | FDR p-value |
| --- | --- | --- | --- |
| *F99 Passage 0* |  |  |  |
| GO:BP | developmental process | GO:0032502 | 4.2E-18 |
| GO:BP | system development | GO:0048731 | 1.3E-16 |
| GO:BP | multicellular organism development | GO:0007275 | 1.1E-15 |
| GO:BP | anatomical structure morphogenesis | GO:0009653 | 3.6E-15 |
| GO:BP | tissue development | GO:0009888 | 2.1E-14 |
| GO:BP | cellular developmental process | GO:0048869 | 7.8E-14 |
| GO:BP | cell differentiation | GO:0030154 | 2.3E-13 |
| GO:BP | cell development | GO:0048468 | 3.6E-09 |
| GO:BP | tube development | GO:0035295 | 3.8E-09 |
| GO:BP | blood vessel development | GO:0001568 | 4.4E-09 |
| GO:BP | circulatory system development | GO:0072359 | 4.6E-09 |
| GO:BP | regulation of cell differentiation | GO:0045595 | 6.0E-09 |
| GO:BP | nervous system development | GO:0007399 | 1.0E-07 |
| GO:BP | epithelium development | GO:0060429 | 3.7E-07 |
| KEGG | TGF-beta signaling pathway | KEGG:04350 | 3.2E-02 |
| WP | miR-509-3p alteration of YAP1/ECM axis | WP:WP3967 | 5.1E-04 |
| WP | Epithelial to mesenchymal transition in colorectal cancer | WP:WP4239 | 2.1E-02 |
| WP | Senescence and Autophagy in Cancer | WP:WP615 | 3.2E-02 |
| *M5 Passage 0* |  |  |  |
| GO:MF | oxidoreductase activity | GO:0016491 | 1.0E-18 |
| GO:MF | electron transfer activity | GO:0009055 | 7.9E-16 |
| GO:MF | proton transmembrane transporter activity | GO:0015078 | 1.3E-12 |
| GO:MF | cytochrome-c oxidase activity | GO:0004129 | 2.3E-07 |
| GO:MF | NADH dehydrogenase activity | GO:0003954 | 3.4E-07 |
| GO:BP | oxidative phosphorylation | GO:0006119 | 4.9E-29 |
| GO:BP | cellular respiration | GO:0045333 | 5.1E-25 |
| GO:BP | lipid oxidation | GO:0034440 | 4.0E-05 |
| GO:BP | fatty acid oxidation | GO:0019395 | 2.1E-04 |
| GO:BP | lipid catabolic process | GO:0016042 | 9.2E-04 |
| GO:BP | fatty acid metabolic process | GO:0006631 | 1.1E-03 |
| GO:CC | mitochondrial part | GO:0044429 | 1.6E-47 |
| GO:CC | mitochondrion | GO:0005739 | 3.8E-37 |
| KEGG | Oxidative phosphorylation | KEGG:00190 | 1.5E-31 |
| KEGG | Citrate cycle (TCA cycle) | KEGG:00020 | 7.0E-05 |
| KEGG | Fatty acid metabolism | KEGG:01212 | 1.6E-03 |
| WP | Electron Transport Chain (OXPHOS system in mitochondria) | WP:WP111 | 2.3E-38 |
| WP | Fatty Acid Beta Oxidation | WP:WP143 | 2.2E-04 |

Note: See Figure S3A for complete set of inter-media comparisons.
FDR, false discovery rate; GO, gene ontology; BP, biological process; MF, molecular function; CC, cellular component; KEGG, Kyoto encyclopedia of genes and genomes; WP, WikiPathways.

### Senescence of primary CEnC is not dependent on mitogen concentration

To determine whether the observed senescent-associated morphogenic changes coincide with gene expression changes, we assessed the expression changes for genes previously associated with cell senescence (Table S6). Many of these genes demonstrated marked expression changes and trends in expression that are consistent with the senescent phenotype observed for many distinct cell systems. To determine the functional consequences of changes in gene expression, we identified genes that were significantly differentially expressed at each passage (P0 was the reference), and for each medium (Table S7). Passage 3 and 4 showed the greatest number of differentially expressed genes (Fig. S3B), which coincided with marked changes in morphometric measures. We subsequently performed GO and pathway analysis to determine the GO and pathway terms enriched in this data set (Table S8). GO terms significantly (q-value<0.05) enriched in the F99 (P3 and P4) differentially expressed genes data set were associated with cell senescence-associated GO terms (e.g., mitotic cell cycle, response to stress and MAPK cascade), and enriched pathway terms were also associated with senescence (e.g., p53 signaling pathway and cellular senescence)(Table 2). Similarly, the significantly enriched GO and pathway terms in the M5 (P3 and P4) differentially expressed genes data set were also associated with cellular senescence, but showed significant enrichment for pathway terms associated with cellular energy processing (e.g., oxidative phosphorylation and the citric acid (TCA) cycle and respiratory electron transport).

**Table 2.**
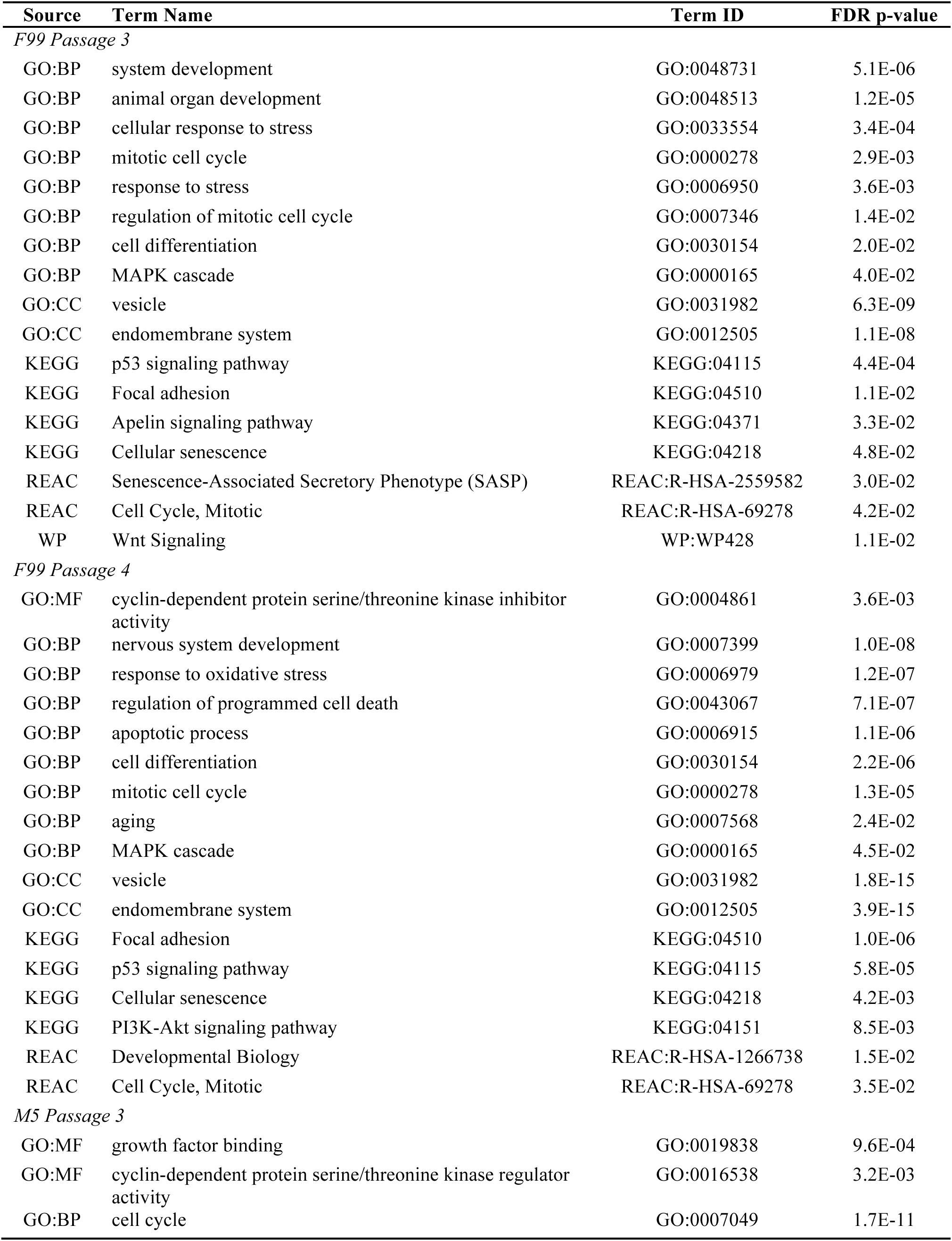
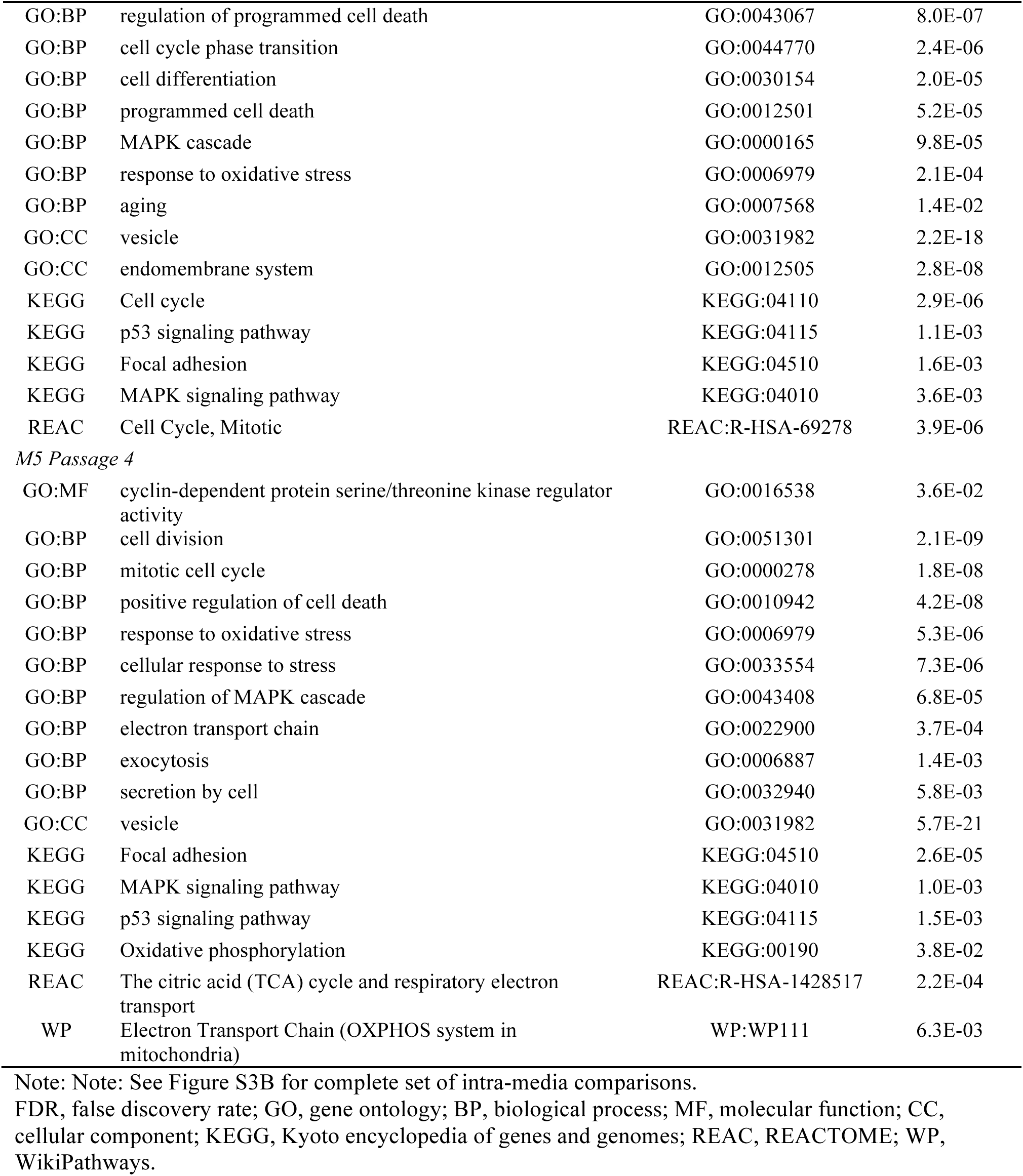
Gene ontology and pathway enrichment analysis of the P3 and P4 intra-media comparisons.

The gene ontology and pathway analyses described above identified biological features in CEnC as a consequence of differentially expressed genes. However, these analyses are performed in the absence of the known direction of differential expression (i.e., downregulated and upregulated) for each gene. To utilize the direction of differential gene expression for the purpose of informing not only pathway enrichment, but also pathway activation state, we used the Ingenuity Pathway Analysis software (Table S9). Significantly activated (z-score>1) pathways in all (P3 (F99 and M5) and P4 (F99 and M5)) data sets were associated with an immune-like response (Table 3). Significantly deactivated (z-score<-1.0) pathways for all data sets were generally associated with metabolic pathways, which included pathways associated with lipid metabolism, nucleotide biosynthesis, glycolysis and cellular respiration.

**Table 3.** Activation state of Ingenuity canonical pathways in P3 and P4 intra-media comparisons.

| **Ingenuity Canonical Pathways** | **z-score** |
| --- | --- |
| *F99 Passage 3* |  |
| NRF2-mediated Oxidative Stress Response | 3.36 |
| IL-8 Signaling | 2.89 |
| Apelin Endothelial Signaling Pathway | 2.71 |
| Cell Cycle: G2/M DNA Damage Checkpoint Regulation | 2.65 |
| Chemokine Signaling | 2.33 |
| IL-3 Signaling | 2.33 |
| ERK/MAPK Signaling | 1.90 |
| VEGF Family Ligand-Receptor Interactions | 1.89 |
| IL-6 Signaling | 1.89 |
| IGF-1 Signaling | 1.67 |
| Acute Phase Response Signaling | 1.67 |
| JAK/Stat Signaling | 1.41 |
| p53 Signaling | 1.13 |
| FGF Signaling | 1.00 |
| Estrogen-mediated S-phase Entry | -1.00 |
| Cyclins and Cell Cycle Regulation | -1.13 |
| RhoGDI Signaling | -1.41 |
| LXR/RXR Activation | -2.24 |
| *F99 Passage 4* |  |
| Cell Cycle: G2/M DNA Damage Checkpoint Regulation | 3.46 |
| NRF2-mediated Oxidative Stress Response | 3.27 |
| IL-8 Signaling | 3.09 |
| Chemokine Signaling | 2.89 |
| p53 Signaling | 2.83 |
| UVC-Induced MAPK Signaling | 2.33 |
| IGF-1 Signaling | 1.41 |
| Inflammasome pathway | 1.34 |
| Apelin Endothelial Signaling Pathway | 1.28 |
| IL-6 Signaling | 1.21 |
| IL-1 Signaling | 1.13 |
| HGF Signaling | 1.00 |
| JAK/Stat Signaling | 1.00 |
| Estrogen-mediated S-phase Entry | -1.13 |
| Chondroitin Sulfate Degradation (Metazoa) | -1.34 |
| Pyrimidine Deoxyribonucleotides De Novo Biosynthesis I | -1.34 |
| NER Pathway | -1.90 |
| LXR/RXR Activation | -3.00 |
| *M5 Passage 3* |  |
| NRF2-mediated Oxidative Stress Response | 3.27 |
| Cell Cycle: G2/M DNA Damage Checkpoint Regulation | 3.16 |
| Chemokine Signaling | 2.71 |
| IL-8 Signaling | 2.69 |
| Acute Phase Response Signaling | 2.50 |
| IL-6 Signaling | 2.32 |
| TGF-β Signaling | 2.11 |
| Inflammasome pathway | 2.00 |
| IL-1 Signaling | 1.89 |
| JAK/Stat Signaling | 1.51 |
| p53 Signaling | 1.39 |
| p38 MAPK Signaling | 1.00 |
| Glycolysis I | -1.00 |
| Fatty Acid β-oxidation I | -1.00 |
| Estrogen Biosynthesis | -1.00 |
| Pyrimidine Deoxyribonucleotides De Novo Biosynthesis I | -2.00 |
| NER Pathway | -2.33 |
| LXR/RXR Activation | -3.00 |
| *M5 Passage 4* |  |
| NRF2-mediated Oxidative Stress Response | 4.60 |
| IL-8 Signaling | 3.51 |
| Chemokine Signaling | 3.00 |
| IL-6 Signaling | 2.84 |
| Cell Cycle: G2/M DNA Damage Checkpoint Regulation | 2.67 |
| Inflammasome pathway | 2.24 |
| p53 Signaling | 2.07 |
| IL-1 Signaling | 1.90 |
| Acute Phase Response Signaling | 1.88 |
| p38 MAPK Signaling | 1.70 |
| JAK/Stat Signaling | 1.41 |
| IGF-1 Signaling | 1.34 |
| Fatty Acid β-oxidation I | -1.63 |
| Acetyl-CoA Biosynthesis I (Pyruvate Dehydrogenase Complex) | -2.00 |
| Cholesterol Biosynthesis I | -2.00 |
| Superpathway of Cholesterol Biosynthesis | -2.45 |
| TCA Cycle II (Eukaryotic) | -2.65 |
| Oxidative Phosphorylation | -5.11 |

Notes: 1) See Figure S3B for complete set of intra-media comparisons. 2) A positive z-score denotes activation and a negative z-score denotes deactivation.

Cell senescence is characterized by an irreversible cell cycle arrest. To determine the impact of passaging (i.e., expansion) on the cell cycle, we examined the intra-media data sets by focusing on the predicted activation state (as predicted by IPA software) of cell cycle functions (Table S10). Generally, this analysis demonstrated activation (z-score>0.5) of cell senescence-associated functions (e.g., senescence of cells) and deactivation (z-score<-1.0) of cell cycle functions that have a positive effect on the cell cycle (e.g., mitosis and cell and cell cycle progression) (Table 4).

**Table 4.** Activation state of cell cycle functions in P0 and P4 intra-media comparisons.

| Function Annotation | z-score |
| --- | --- |
| *F99 Passage 3* |  |
| Senescence of cells | 0.99 |
| Replicative senescence of cells | 0.60 |
| Cell cycle progression | -0.86 |
| Re-entry into interphase | -1.34 |
| Mitosis | -1.72 |
| Initiation of interphase | -1.73 |
| *F99 Passage 4* |  |
| Senescence of tumor cell lines | 1.51 |
| G1 phase | 1.35 |
| Senescence of cells | 0.69 |
| M phase | -0.53 |
| M phase of cervical cancer cell lines | -0.79 |
| M phase of tumor cell lines | -1.30 |
| *M5 Passage 3* |  |
| Senescence of cells | 1.14 |
| G1 phase | 0.90 |
| Cell cycle progression | -0.81 |
| M phase | -1.19 |
| Mitosis | -1.69 |
| Segregation of chromosomes | -2.13 |
| *M5 Passage 4* |  |
| G1 phase | 2.24 |
| Senescence of cells | 1.70 |
| Cell cycle progression | -0.88 |
| Cytokinesis | -1.06 |
| M phase | -1.18 |
| Segregation of chromosomes | -1.69 |

Notes: 1) See Figure S3B for complete set of intra-media comparisons. 2) A positive z-score denotes activation and a negative z-score denotes deactivation.

### Senescence of primary CEnC involves the activation of the p53 and p38-MAPK pathways

To determine whether senescence of primary CEnC involves activation of the p53 and p38-MAPK pathways and regulation of the cell cycle, we examined expression and activation of genes/proteins known to regulate cell cycle arrest due to senescence (Fig. 4). We created a custom pathway network that included p53 (TP53) and p38-MAPK (MAPK14), the cyclin-dependent kinase inhibitors p21^CIP1^ (CDKN1A) and p16^INK4^ (CDKN2A), and well-established markers of cell senescence (LMNB1, GADD45A and CD44) (Fig. 4A). In addition, we included the “senescence of cells” as the phenotype connecting these factors. Initially, we superimposed the differential gene expression results (fold-change) for genes encoding these proteins, and then we applied an algorithm that predicted the “activation” state of the other proteins and the phenotype. Applying the prediction algorithm demonstrated activation of both p53 and p38-MAPK, and activation of “senescence of cells” phenotype in primary CEnC at P3 in both F99 and M5. Detection of phosphorylation of p53 and p38-MAPK at P3 by Western blotting provided confirmation of their activation (Fig. 4B). Although *TP53* and *MAPK14* gene expression were not markedly different between P0 and P3 CEnC, phosphorylation of the encoded proteins was significantly greater at P3 than P0 (p<0.05)(Fig. 4C). In addition, significant (p<0.05) differential gene expression of the senescence markers *LMNB1*, *GADD45A* and *CD44* was observed at P3 in either F99 or M5 or both, a result consistent with cell senescence (Fig. 4D). In addition, we confirmed significantly increased expression of the CD44 protein at P3, which was consistent with *CD44* transcript levels.

**Figure 4.**
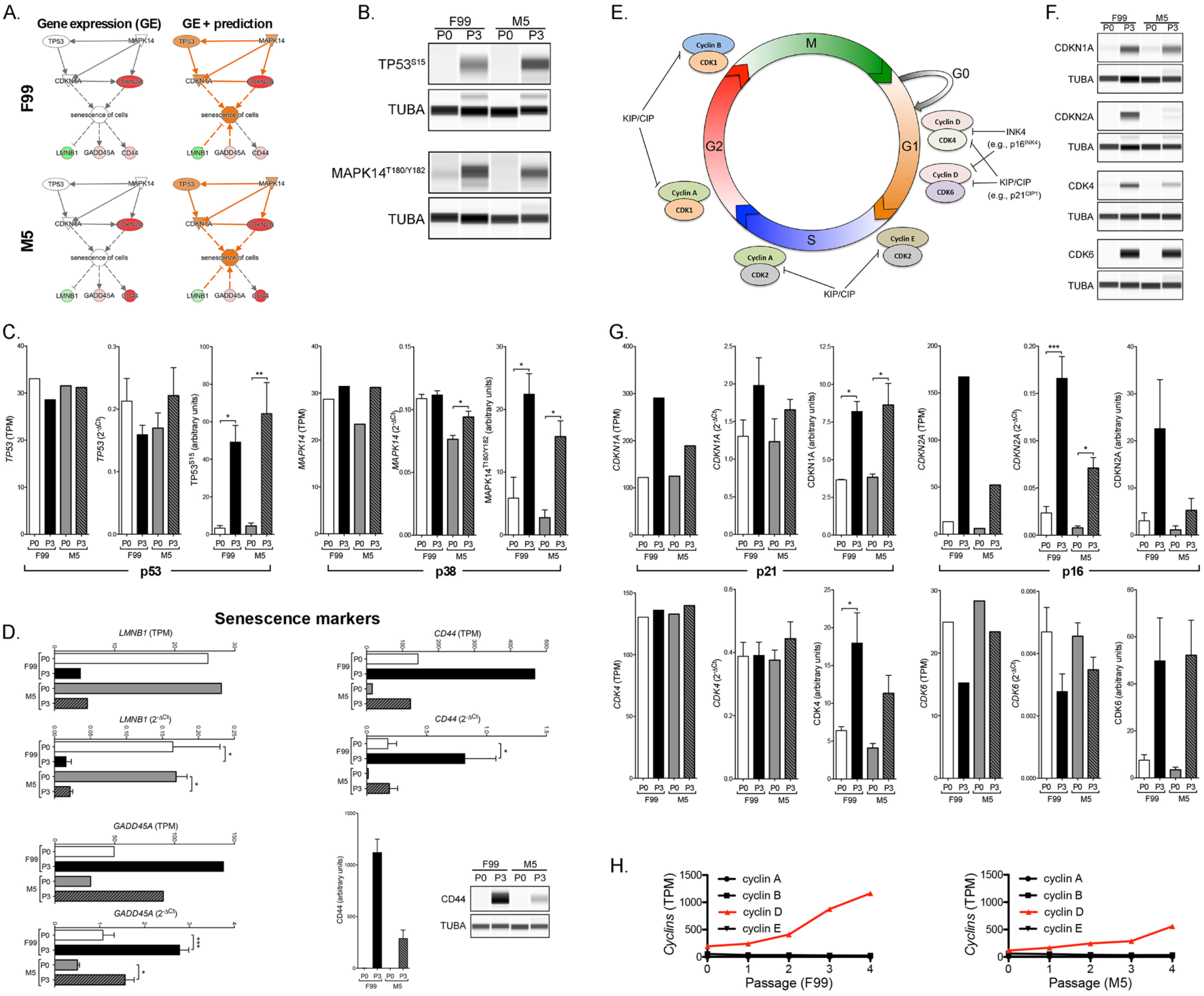
Cellular senescence of cultured CEnC is associated with activation of p53 and p38-MAPK pathways. (A) Classic cell senescence pathway involving p53 (TP53) and p38 (MAPK14). Pathway schematics show expression of senescence-associated genes that were differentially expressed in P3 CEnC (left panel). Shades of green, decreased expression; shades of red, increased expression. Pathway schematics show differentially expressed genes together with activity prediction results (orange, activated; blue, repressed) for upstream and downstream factors (right panel). (B) Western results show phosphorylation of TP53 at Serine 15 and MAPK14 at Threonine 180 and Tyrosine 182 in CEnC at P0 and P3, cultured in either F99 or M5. (C) Bar graphs show *TP53* and *MAPK14* transcript levels in P0 and P3 CEnC using RNA-seq and validated by qPCR. Bar graphs show the quantification of TP53 and MAPK14 phosphorylation in P0 and P3 CEnC. (D) Bar graphs show the transcript abundances of *LMNB1*, *GADD45A* and *CD44*, and validated using qPCR. Bar graphs show CD44 protein abundances detected using a Western assay. (E) Diagram of the cell cycle including factors that positively (cyclins and CDK) and negatively (INK4 and KIP/CIP) regulate cell cycle progression. (F) The p21^CIP1^ (CDKN1A) and p 16^INK4^ (CDKN2A) are inhibitors of cyclin-dependent kinases, and are classically associated with cellular senescence. Western results show increased expression of CDKN1A and CDKN2A concomitant with increased expression of the CDK4 and CDK6 in CEnC at P3, cultured in either F99 or M5. (G) Bar graphs show: *CDKN1A*, *CDKN2A*, *CDK4* and *CDK6* transcript levels in P0 and P3 CEnC using RNA-seq and validated by qPCR; the quantification of the proteins encoded by these genes in P0 and P3 CEnC. (H) Line graphs show the transcript abundances of the cyclins at each passage in CEnC cultured in F99 or M5. TUBA was used as a loading control in the Westerns. Data in bar graphs are represented as the mean ±SEM (n=4, mRNA; n=3, protein). Statistical comparisons were performed using one-way ANOVA with post-hoc Tukey test. *, P<0.05; **, P<0.01; ***, P<0.001.

The cell cycle includes various phases (G1, S, G2 and M) that involve cyclin and cyclin-dependent kinases (CDK) (Fig. 4E). In the absence of inhibitors of the CDK, cell cycle progression is dependent on the availability and interaction of cyclins and CDK. However, two classes of inhibitors, INK4 and KIP/CIP, are able to inhibit cell cycle progression via interaction of the inhibitors (e.g., p21^CIP1^ and/or p16^INK4^) with the cyclin/CDK complexes, an interaction that ultimately blocks the complexes’ downstream actions and results in cell cycle arrest. We assessed protein expression of both p21^CIP1^ (CDKN1A) and p16^INK4^ (CDKN2A), which are upregulated in senescent cells (Fig. 4F). Marked increases in both of these inhibitors were observed in P3 CEnC cultured in F99 or M5. In addition, accumulation of CDK4 and CDK6, CDK important in progression through G1/S of the cell cycle, was observed in P3 cells cultured in F99 and M5 (Fig. 4F). While transcript quantification demonstrated a significant increase in gene expression in P3 cells only for CDKN2A, the expression of the encoded proteins were either significantly (p<0.05; CDKN1A, CDK4) or markedly (CDKN2A, CDK4 and CDK6) increased in P3 cells cultured in F99 and M5 (Fig. 4G)., As an increase in cyclin D, in conjunction with increases in p16^INK^ (CDKN2A), CDK4 and CDK6, is a hallmark of G1 cell cycle arrest, we measured the expression of the four cyclins (cyclins A, B, D and E). A marked and progressive increase in cyclin D was observed in cells cultured in both F99 and M5, in contrast to the expression of the other three cyclins, which was unchanged through P4 (Fig. 4H).

### Low mitogenic conditions establish a robust functional barrier in CEnC

To determine the impact of cell culture conditions on the establishment of a functional barrier, we measured resistance of the CEnC to an electrical current (Fig. 5). Initially, we measured barrier function of cells isolated/grown using unmodified protocols for each of the two methods employed in this study (Tryp-LN-F99 and ColA-ColIV-M4M5) CEnC cultured using the ColA-ColIV-M4M5 method demonstrated markedly greater impedance compared with CEnC cultured using the Tryp-LN-F99 method (Fig. 5A). Impedance data was then used to model cell-cell (*R_b_*) and cell-substrate (*α*) interactions, which demonstrated that both contribute to the greater barrier function observed in cells cultured using the ColA-ColIV-M4M5 method. To identify a potential effect of the dissociation enzymes (trypsin and collagenase A) on the establishment of a functional barrier, we modified the isolation/growth protocols to exchange one enzyme for the other. Exchanging the enzymes did not have a marked effect on the establishment of a functional barrier, with the cells cultured using the protocol containing ColIV-M4M5 establishing a more robust functional barrier compared with cultures established with the protocol containing LN-F99 (Fig. 5B). To determine the contribution, if any, of the substrate (laminin or collagen) to which the cells adhere, or of the media (F99 or M5) in which the cells are cultured, we exchanged the substrates for each other and used F99 and M5 for each (Fig. 5C). Due to the limitation in the number of replicates we could assay, we chose trypsin (Tryp) as the dissociation enzyme, since use of either trypsin or collagenase A did not make a significant difference on establishment of a functional barrier. While only marginal differences in the electrical impedance were observed up to approximately 48 hours, after that time point the cells that were cultured using the Tryp-LN-M4M5 method demonstrated a progressively weakening barrier. Additionally, the cells that were cultured using the Tryp-LN-M4M5 method demonstrated a marked decrease in cell-cell and cell-substrate adhesion after 48 hours.

**Figure 5.**
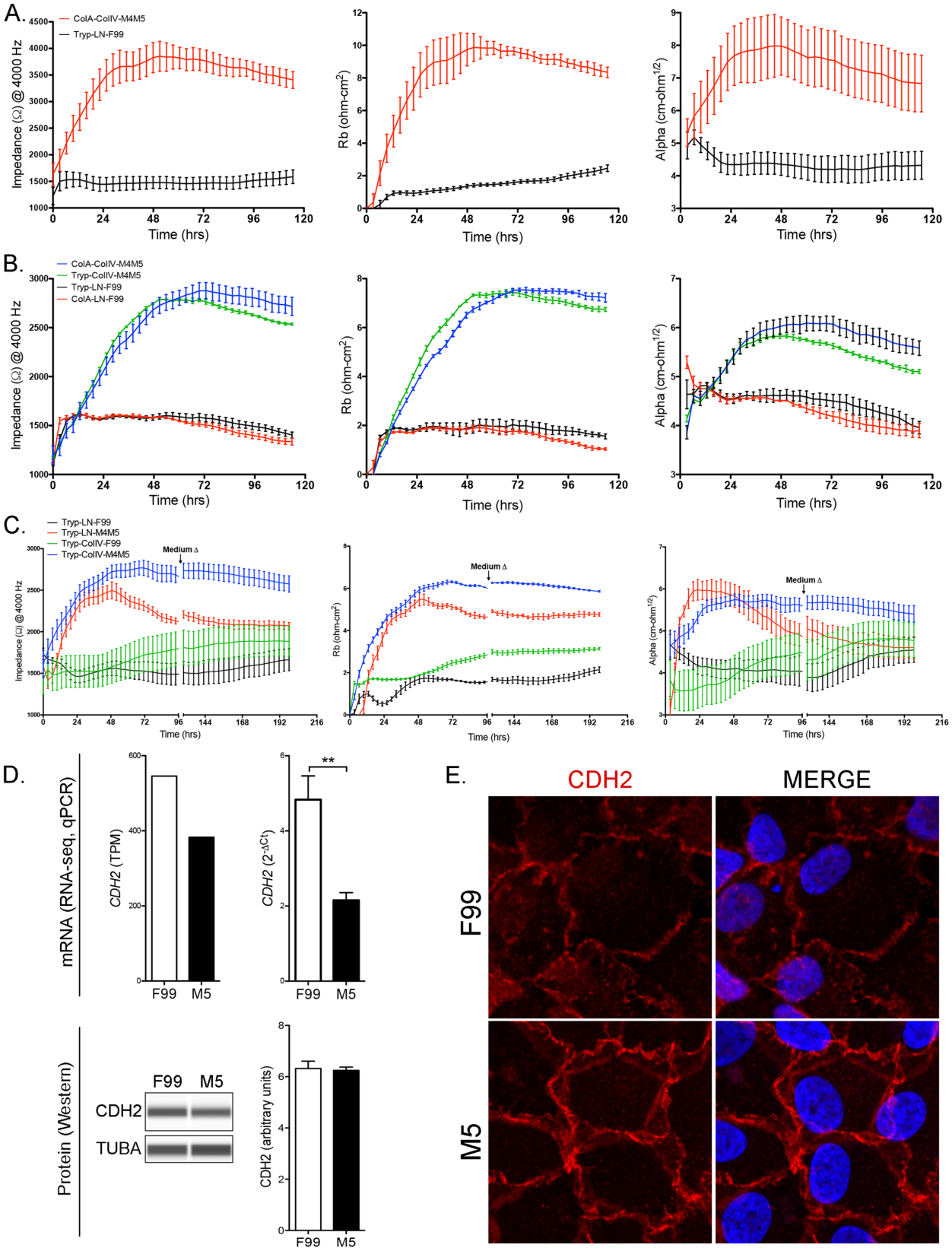
Cultured CEnC establish and maintain robust barrier function in M5 medium. (A) Line graph shows electrical impedance (Ω at 4000 Hz) for P0 CEnC monolayers established in F99 and M5 media. The contribution of cell-cell adhesion to the resistance of electrical current was modeled from impedance data and given as *R*_b_ (Ω • cm^2^). The contribution of cell-substrate adhesion to resistance of electrical current was modeled from impedance data and given as (α, Ω^½^ • cm). (B) Line graph shows electrical impedance for CEnC monolayers established from cells dissociated with trypsin (Tryp) or collagenase (ColA), and each cultured in F99 and M5 medium. The contribution of cell-cell (*R*_b_) and cell-substrate (α) to electrical resistance were modeled from the impedance data. (C) Line graph shows electrical impedance for CEnC monolayers established from cells dissociated with trypsin (Tryp), seeded on either laminin (LN) or collagen (ColIV), and cultured in either F99 and M5 medium. The contribution of cell-cell (*R*_b_) and cell-substrate (*α*) to electrical resistance was modeled from the impedance data. (D) Bar graphs show abundance of *CDH2*, a contributor to cell-cell adhesion, in CEnC cultured in F99 and M5 (top panel). Western results for CDH2 protein and bar graph shows protein quantification (bottom panel). TUBA was used as a loading control in the Western. (E) Immunofluorescence results show localization of CDH2 protein in CEnC cultured in F99 and M5. Red, Alexafluor 594; blue, DAPI. Data in line graphs are represented as the mean ±SD (n=4), and data in bar graphs are represented as the mean ±SEM (n=4, mRNA; n=3, protein). Statistical comparisons were performed using one-way ANOVA with post-hoc Tukey test. **, P<0.01.

To determine whether the differences observed in barrier function between the two culturing methods (F99 versus M5) could be explained by differences in barrier-associated proteins or in the organization of the adhesive interactions between cells, we examined the expression and localization of the cadherin protein CDH2. While *CDH2* transcript levels measured by qPCR were significantly (p<0.01) lower in cells cultured in M5, the CDH2 protein levels were not different (Fig. 5D). However, the organization of the adhesive interaction between cells was markedly different between cells grown in F99 and M5 (Fig. 5E). In F99, the cell-cell interactions appeared diffuse and frayed, although these features were not consistent for all cell-cell interactions, and lacked features of a well-formed lateral membrane. In contrast, the cells grown in M5 appeared to have consistent cell-cell interactions with what appears as prominent lateral membranes, which give the image the impression of depth.

### A strong functional barrier in primary CEnC is associated with robust expression of adhesion and glycocalyx proteins

To determine the extent to which barrier-associated proteins may be involved in the more robust establishment of a functional barrier of CEnC in M5 media, we examined several proteins associated with either cell-cell adhesion or glycocalyx formation (Fig. 6). We first measured the expression of genes that encode adhesion-associated proteins (CLDN11, AJAP1, TMEM204 and TMEM178A), and quantified protein levels for each (Fig. 6A). The expression of each of the four genes was higher in CEnC cultured in M5 compared to F99, reaching statistical significance (p<0.05) for all four when measured at the transcript level by qPCR, and for two (CLDN11 and TMEM178A) when measured at the protein level (p<0.01). We then measured the expression of genes that encode glycocalyx-associated proteins (APOE, MYOC, DCN, LUM and APOD), and quantified protein levels for each (Fig. 6B). Significantly (p<0.05) higher transcript levels were observed for all five genes in CEnC cultured in M5, but only two of the proteins (MYOC and APOD) demonstrated significantly higher levels. The APOE (p=0.05) and LUM (p=0.14) proteins were markedly higher in M5, but did not achieve statistical significance. DCN did not demonstrate a difference in protein levels between the two media. Of note, APOD is known to exist in three forms (pre-modified, post-modified, and complexed), each of progressively greater molecular weight, all of which demonstrated significantly (p<0.05) higher expression in M5 compared with F99.

**Figure 6.**
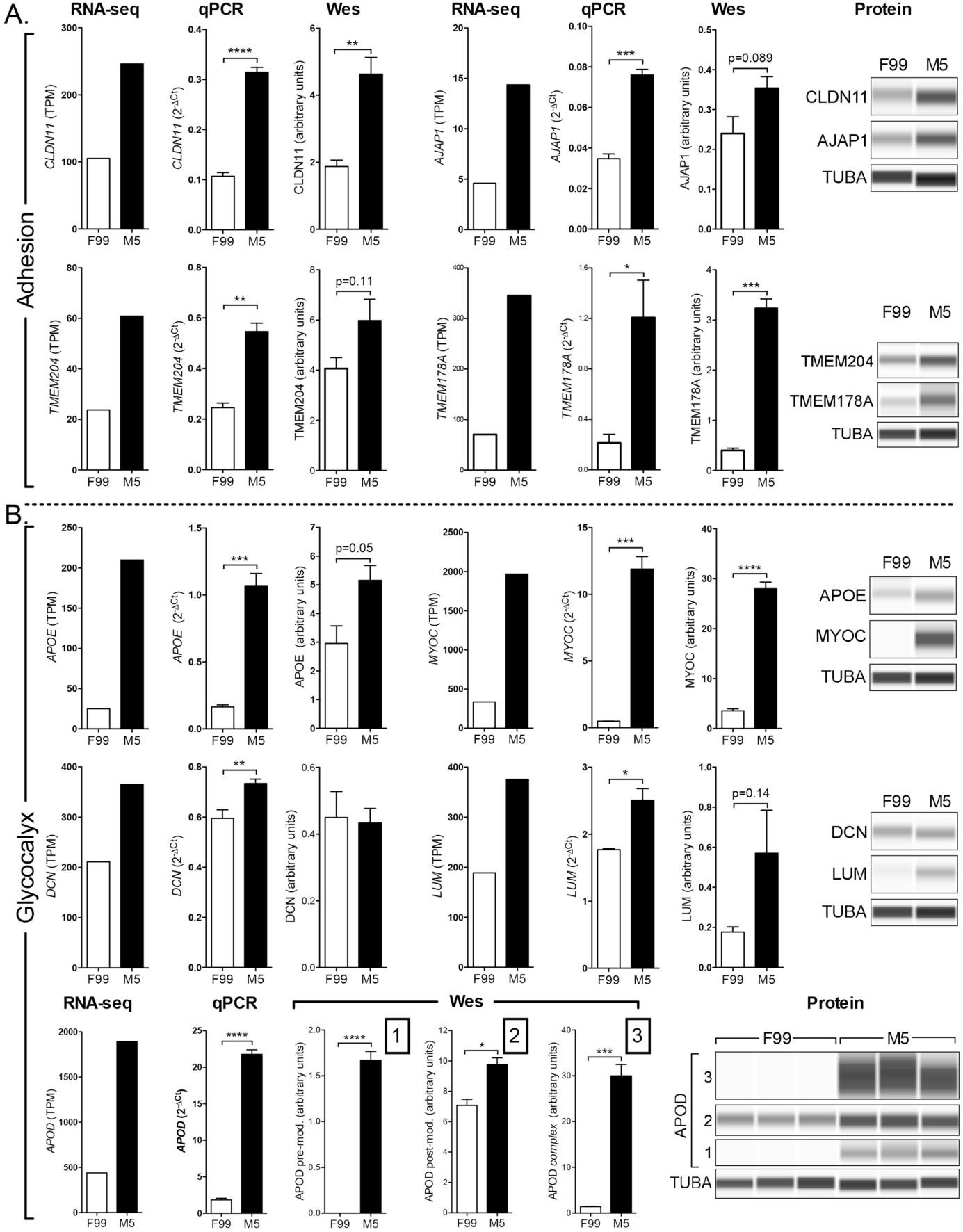
M5 medium increases the expression of barrier-associated genes and proteins. (A) Bar graphs show the expression levels (RNA-seq and qPCR) of genes associated with cell adhesion (*CLDN11*, *AJAP1*, *TMEM204* and *TMEM178A*) in P0 CEnC. Detection and quantification of the proteins encoded by these genes were performed using a Western assay (Wes). (B) Bar graphs show the expression levels (RNA-seq and qPCR) of genes associated with the glycocalyx (*APOE*, *MYOC*, *DCN*, *LUM and APOD*). Detection and quantification of the proteins encoded by these genes were performed using a Western assay (Wes). Three forms of APOD were observed, and each was quantified and graphed. The protein forms observed were a pre-modified form (1), a post-modified form (2) and a high molecular weight complex form (3). TUBA was used as a loading control in the Westerns. Data in bar graphs are represented as the mean ±SEM (n=4, mRNA; n=3, protein). Statistical comparisons were performed using one-way ANOVA with post-hoc Tukey test. *, P<0.05; **, P<0.01; ***, P<0.001; ****, P<0.0001.

### Robust CEnC pump function is established by culture in low mitogenic conditions

The SLC4A4, SLC4A11 and SLC16A1 transporters are highly expressed in in vivo corneal endothelium, and play a prominent role in maintaining the solute gradients necessary for regulation of water transport from the stroma to the anterior chamber. To determine the level of expression and functional capacity of each in F99 and M5 media, we measured transcript levels by RNA-seq and qPCR, protein levels by Western, and transporter activity with a fluorescence-based transporter assay (Fig. 7). *SLC4A11* demonstrated significantly higher (p<0.01) and *SLC4A4* demonstrated non-significantly (p=0.081) higher transcript levels in M5 medium, while the proteins encoded by all three genes demonstrated significantly (p<0.05) higher levels in M5 compared with F99 (Fig. 7A).

**Figure 7.**
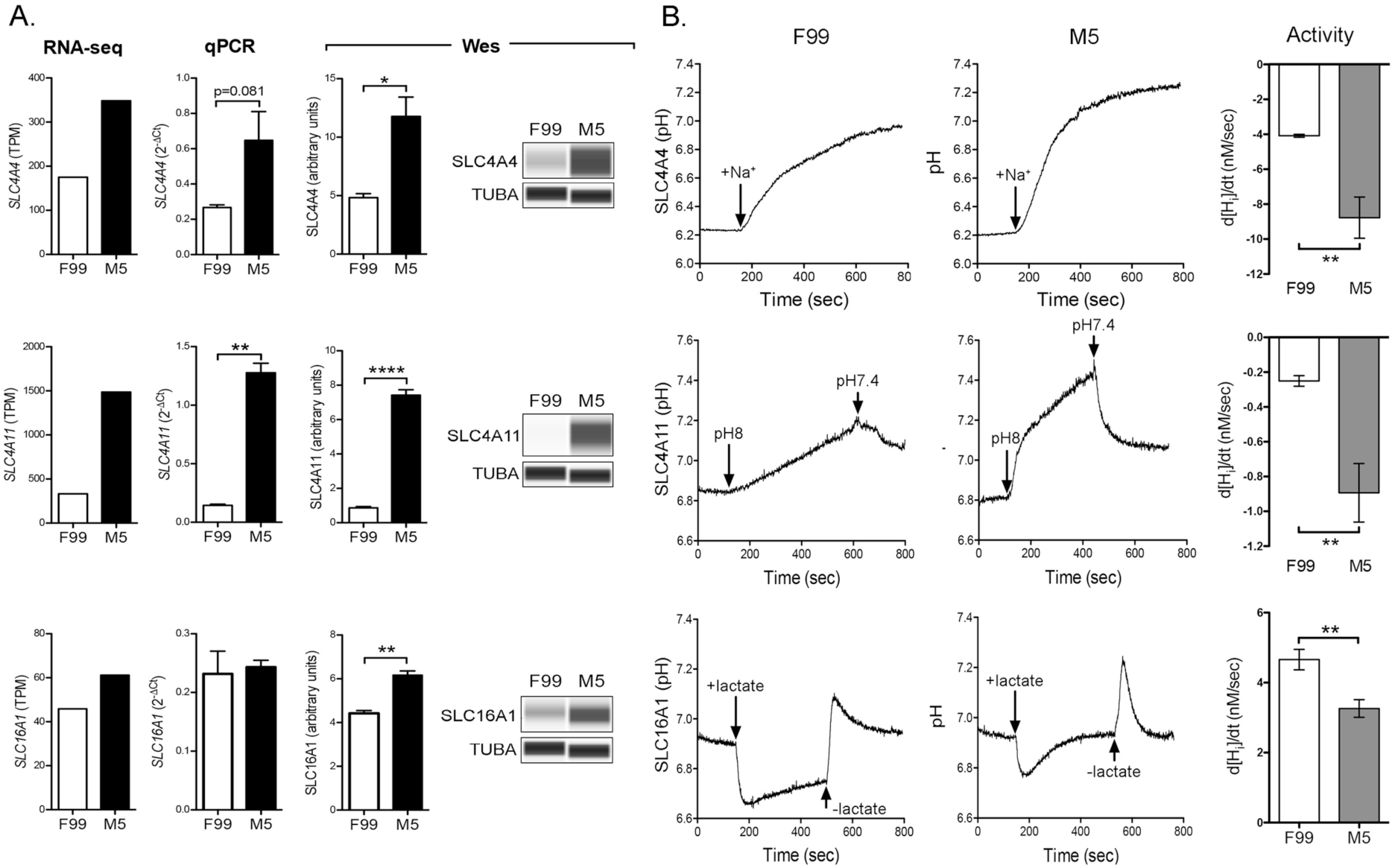
SLC4A4 and SLC4A11 transporter activity is increased in M5 medium. (A) Bar graphs show transcript abundances (RNA-seq and qPCR) in P0 CEnC for genes that encode membrane bound transporters (*SLC4A4*, *SLC4A11* and *SLC16A1*). Detection and quantification of the encoded proteins was performed using a Western assay (Wes). (B) Traces show change in pH levels before and after perfusion with buffer containing indicated substrate specified for the transporter (SLC4A4, Na^+^; SLC4A11, H^+^; SLC16A1, lactate). Arrows indicate time at which buffer was changed. Bar graphs show the rate of change in intracellular proton concentration [Hi], which was calculated as a proxy for transporter activity. Data in bar graphs are represented as the mean ±SEM (n=4, mRNA; n=3, protein; n=6, transporter activity). Statistical comparisons were performed using one-way ANOVA with post-hoc Tukey test. *, P<0.05; **, P<0.01; ****, P<0.0001.

To assess transporter function, we assayed for intracellular proton concentration (i.e., pH_i_) over time, including after changes in buffer perfusion, since each of the transporters transfer protons across the plasma membrane (Fig. 7B). Traces showing intracellular pH demonstrate sensitivity to buffer changes for each transporter. Qualitatively, the rate of change in intracellular proton concentration mediated by SLC4A4 and SLC4A11 following a change in buffer appears greater in M5 compared with F99 medium. In contrast, the rate of change in intracellular proton concentration mediated by SLC16A1 following a change in buffer appears greater in F99 compared with M5 medium. Quantitatively, and as an indirect measure of transporter activity, we calculated the change in intracellular proton concentration over time (d[H_i_]/dt) for each of the assays, and observed significantly (p<0.01) higher SLC4A4 and SLC4A11 and significantly (p<0.01) lower activity for SLC16A1 for cells cultured in M5.

### Low mitogenic conditions cause decreased cell migration capacity

We measured cell migration using a non-wounding method (Fig. 8). Phase contrast imaging of cell migration demonstrated significantly (p<0.001) less gap closure for CEnC in M5 compared with F99 medium (Fig. 8A and B). We observed that cells cultured in M5, while possessing lower migration capacity, also appear to markedly cover the gap primarily by increasing the cell size rather than by migration (Fig. 8C). We calculated the ratio of the area occupied by the cells that “migrated” into the gap versus cells more distant from the gap, and which have not substantially changed their size compared with prior to when migration was initiated (Fig. 8D). Cell size for cells cultured in M5 and occupying any portion of the gap cover a significantly (p<0.05) larger area compared with cells cultured in F99.

**Figure 8.**
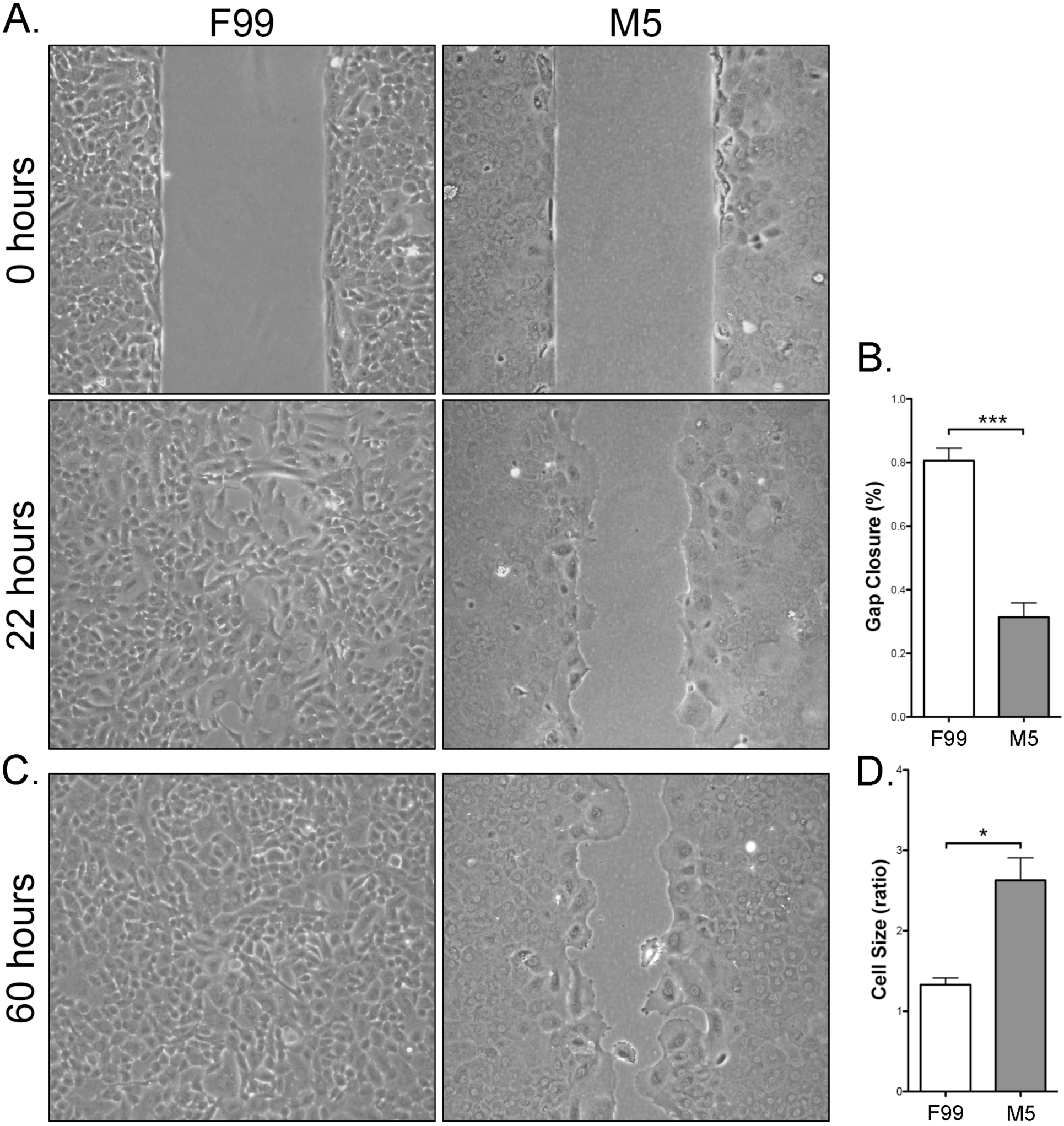
CEnC cultured in M5 medium possess attenuated migration capacity. (A) Phase contrast images show initial gap (0 hours) and gap closure (22 hours) of P0 CEnC cultured in F99 and M5. (B) Bar graph shows quantification of gap closure at 22 hours as a percent of the initial gap. (C) Phase contrast images show gap closure at 60 hours. (D) Bar graph shows the ratio of the mean area at 60 hours of the cells near or populating the gap divided by the mean area of the cells not near the gap. Data in bar graphs are represented as the mean ±SEM (n=4). Statistical comparisons were performed using one-way ANOVA with post-hoc Tukey test. *, P<0.05; ***, P<0.001.

## DISCUSSION

Corneal clarity depends on the corneal endothelium, and disease or injury to this tissue causes loss of visual acuity that may necessitate corneal transplantation. The development and widespread adoption of endothelial keratoplasty techniques has significantly improved outcomes of the surgical management of corneal endothelial cell dysfunction although postoperative complications remain (Deng et al., 2015; Lass et al., 2017; Van den Bogerd et al., 2018). However, the current paradigm of one donor cornea for one recipient requires cornea donation, recovery and utilization rates that are present in only a few countries in the world that are able to meet the domestic need for transplantable corneas. Thus, scientists and clinicians have developed various methods for the isolation and in vitro culturing of CEnC, with the aim of achieving sufficient cell numbers of transplantable high quality CEnC to challenge the one donor-one recipient paradigm (Soh et al., 2017). While several metrics have been used to assess the identity (e.g., morphology, biomarker expression) and function (e.g., transcellular resistance, intracellular proton concentration) of cultured CEnC, a comprehensive study of the effects of in vitro expansion on the CEnC transcriptome has not been performed. The transcriptomic work performed as part of this study may be used as one of the criteria for assessing the quality and viability of in vitro expanded CEnC for use in cell-replacement therapy. In particular, because cell senescence is a feature of primary cell culture, it is necessary to identify the potential signaling and metabolic pathways involved in CEnC senescence prior to developing strategies to target these pathways and delay senescence, thereby producing a larger number of therapeutically suitable cells.

In 1978, the first human CEnC cultures were described, but given a lack of human-specific protocols, the researchers used culture conditions optimized for growth of rabbit CEnC (Jumblatt et al., 1978). Over the next decade, techniques for the isolation of human CEnC and growth media for the in vitro culture of human CEnC were optimized. In 1989, Engelmann et al. determined that a 1:1 mixture of M199 and F12 media was the optimal base medium for culture of primary CEnC (Engelmann and Friedl, 1989). Two decades later, the formulation of the medium was modified to induce cell cycle re-entry (due to relatively mitogen-rich conditions) and establishment of contact inhibition, leading to quiescence, a non-proliferative cell state that characterize CEnC in vivo (Valtink et al., 2008). We started culturing primary CEnC in 2011, and chose the medium formulation described by Valtink et al. as it was at the time the most effective formulation for the culture of both primary and immortalized CEnC. Nevertheless, about 40% (3/7 at P0 in this study) of the cultures we established did not possess CEnC-like morphology, and were generally characterized by a mixture of morphologies (some resembling fibroblasts), an observation consistent with previous findings by the developers of the formulation (R. Faragher, personal communication, March 2017). In 2015, Peh et al. described a novel approach to in vitro culturing of CEnC in which cell proliferation was accomplished by growth in a mitogenic-rich medium, followed by maintenance in a medium with reduced mitogens (Peh et al., 2015). Coincidently, a recent report independently demonstrated the superiority of the dual media approach (Bartakova et al., 2018). Taken together, these studies indicated that the low mitogenic medium established cultures with better morphology, marker expression and function compared with growth in a mitogen-rich medium alone. Herein, we compared the two culture methods published by Valtnik et al. and Peh et al. using extensive bioinformatics analyses and additional functional techniques to determine which method performs best in maintaining the CEnC phenotype throughout in vitro expansion.

Cell morphology is an initial metric for assessing the quality of CEnC cultures. We observed that the low mitogenic environment was better at maintaining a CEnC-like morphology, but continued passaging caused a distinct morphogenic transformation consistent with cellular senescence. This change in morphology correlated with a progressive loss of CEnC identity as defined by the expression of evCEnC-specific genes. Temporal changes in evCEnC-specific gene expression as a consequence of passaging allowed us to identify a novel cell surface marker, encoded by *TMEM178A*, as a potential positive selection marker for high quality cultured CEnC. In addition, after a reassessment of reported selection markers, we propose SLC4A11 (positive), TMEM178A (positive) and CD44 (negative) as the minimum for selection of high quality in vitro CEnC (Bartakova et al., 2018; Cheong et al., 2013; Chng et al., 2013; Ding et al., 2014; Frausto et al., 2016; Okumura et al., 2014a; Toda et al., 2017; Yoshihara et al., 2015). SLC4A11 is a highly expressed transporter protein in the corneal endothelium that plays a significant role in CEnC function, and biallelic mutation of this protein leads to congenital hereditary endothelial dystrophy (Aldave et al., 2013; Jiao et al., 2007; Vithana et al., 2006). Conversely, CD44 is not expressed in the corneal endothelium, and its expression correlates with a senescent phenotype in vascular and corneal endothelial cells (Mun and Boo, 2010; Ueno et al., 2016).

A prominent barrier to expansion of CEnC in vitro for use in a cell-based therapy is the limited proliferative capacity of primary CEnC. In vivo, CEnC are in a state of cellular quiescence, which in this cell type is specifically a cell cycle arrest at the G1 phase that is established by contact-inhibition, maintained by TGFβ2, and regulated by p27^KIP1^ (Joyce, 2003). However, in vivo CEnC have been shown to express senescent markers in older donors, which is postulated to contribute to the lower proliferative capacity of primary CEnC isolated from older donors compared with those from younger donors. Nevertheless, CEnC are able to re-engage the cell cycle by loosening of cell-cell contacts and/or exposure to potent mitogenic factors. As with most primary cells, primary CEnC have a limited life span under proliferative (i.e., passaging and growth in mitogen-rich) conditions. This limitation is caused by cellular senescence of CEnC (in vitro and ex vivo), and may be, in part, a consequence of oxidative stress (Joyce et al., 2009). This is supported by our observations in late passage CEnC (P3 and/or P4) that p38MAPK was phosphorylated (stress response), that expression of p16^INK4^ (gene and protein) was increased, and that the set of differentially expressed genes was enriched for genes implicated in oxidative stress-associated pathways (e.g., NRF2-mediated oxidative stress response, DNA damage checkpoint regulation). In addition, there is also an apparent repression of oxidative phosphorylation (i.e., mitochondrial activity), which was observed in our pathway analysis. Moreover, we also identified enrichment for genes associated with p53 signaling, an observation made previously (Sheerin et al., 2012), and this coincided with phosphorylation of p53 and increased expression of p21^CIP1^ (CDKN1A) (gene and protein) in senescent CEnC (i.e., P3 and P4). Cell senescence in primary CEnC in this study was also associated with increased expression of CDK4 and CDK6 proteins (but not gene expression), and increased *cyclin D* (*CCND1*) gene expression. Taken together, this represents a gene/protein expression profile consistent with senescence-induced G1 cell cycle arrest. Overall, activation of the p38MAPK and p53 pathways did not appear significantly influenced by growth in a low- or high-mitogenic environment.

Achieving and maintaining characteristic CEnC functional properties is essential for the clinical utility of in vitro CEnC in management of CEnC dysfunction. The corneal endothelium is a monolayer mosaic of CEnC, and its primary function is to maintain the corneal stroma in a partially dehydrated state to preserve its optical clarity. It does this by establishing a robust barrier conferred by cell-cell and cell-substrate interactions, which allow for passive leak of water into the stroma, but not most solutes (Srinivas, 2010). Through the combined action of various solute transporters, unidirectional translocation of water from the stroma into the aqueous humor occurs (Bonanno, 2012). Taken together, this describes the “pump-leak” hypothesis of water homeostasis in the cornea. We demonstrated that a low mitogenic environment combined with culture on collagen represented better in vitro conditions for establishing a robust CEnC barrier (as measured by electrical impedance) compared with a high mitogenic environment and culture on laminin. Recent studies showed that collagen, compared with various other extracellular matrix proteins, including laminin, and low mitogen-containing media were optimal for establishing robust barrier function in a CEnC line (Santander-Garcia et al., 2016) and primary CEnC (Bartakova et al., 2018). Detection of CDH2, a CEnC-specific marker, in CEnC cultures in low mitogenic conditions demonstrated that cell adhesions (cell-cell and cell-substrate) are considerably more complex, and showed striking resemblance to the organization of adhesions in ex vivo human corneas (He et al., 2016). We also demonstrated that this increase in barrier function is associated with an increase in adhesion/glycocalyx-associated protein expression. To our knowledge, this represents the first report implicating glycocalyx proteins in corneal endothelial biology. Although some of the barrier-associated proteins we examined in this study have been associated with the glycocalyx of the vascular endothelium (Friden et al., 2011; Reitsma et al., 2007), it remains to be determined if the increase in glycocalyx-associated proteins coincides with the formation of a glycocalyx in corneal endothelium either in vitro or in vivo. However, early reports suggest that the human corneal endothelium may possess a glycocalyx-like layer composed polysaccharides, glycosaminoglycans and other, as yet, unidentified molecules (Hornung and Wollensak, 1979; Jacobsen and Sperling, 1978; Schroder and Sperling, 1977; Sperling and Jacobsen, 1980). We also showed that at least two transporters critical for CEnC function (SLC4A4 and SLC4A11) are expressed at significantly higher levels and show increased transporter activity in low mitogenic conditions compared with high mitogenic conditions. As such, CEnC expanded in mitogenic-rich medium and maintained in low mitogenic conditions show functional features consistent with the CEnC requirement for regulating water homeostasis in the cornea. These results are consistent with previous studies demonstrating an intact pump function (i.e., prevention/mitigation of edema) in in vitro expanded corneal endothelial cells in an animal model of bullous keratopathy (Peh et al., 2017; Peh et al., 2019).

A long known characteristic of in vivo CEnC is that when mild cell loss occurs, the vacated area is covered by migration and increase in size of adjacent cells (Sherrard, 1976). To determine if this phenomenon is recapitulated in vitro, we performed a non-wounding cell migration assay. Cells in a high mitogenic environment migrated and/or proliferated (not distinguished in our study) to cover the gap with no significant increase in cell size. In contrast, CEnC in low mitogenic conditions behaved in a manner consistent with their behavior in vivo, so that these cells showed low migration capacity, but a significant propensity to increase cell size to cover the gap. In addition, these results are consistent with the differences observed for cell barrier formation, because increased cell-cell adhesion is antithetical to robust cell migration (and also to robust cell proliferation capacity).

In summary, we provide evidence that CEnC proliferation in a high mitogenic medium, followed by maintenance of contact-inhibition of confluent monolayers in a low mitogenic medium, supports establishment of CEnC in which in vivo function is significantly recapitulated and permits sufficient expansion of CEnC to significantly challenge the “one donor-one recipient” paradigm. In our current study, we can theoretically achieve a 1:5-1:15 donor:recipient ratio (donor:recipient, 1:5 (P0) or 1:15 (P1) – at approximately 3000 cells/mm^2^), as these are the passages at which the cells possess the most robust CEnC phenotype as determined by gene (or marker) expression and cell morphology. Maintaining the CEnC morphologic and functional phenotype at higher passage numbers remains an active area of research. Also, while we examined cell function at an early passage (typically at P0), we recognize that functional analysis of cells will need to be performed at later passages to ensure viability as a therapeutic modality. The progression to senescence in vitro remains a significant barrier to the expansion of cultured CEnC. Identification of oxidative stress as a component of CEnC senescence has prompted investigators to add ascorbic acid as a supplement to early media formulations (added to F99 and M4) to reduce/inhibit oxidant-induced stress/apoptosis (Serbecic and Beutelspacher, 2005; Shima et al., 2011) and to use p38MAPK inhibitors to delay the onset of senescence, but this approach has shown mixed results (Hongo et al., 2017; Nakahara et al., 2018; Sheerin et al., 2012). In addition, senescent CEnC in vitro are characterized by the Senescence-Associated Secretory Phenotype (SASP), which includes the secretion of immune-regulating factors (cytokines, chemokines and growth factors) that have been demonstrated to support tumorigenesis in adjacent epithelial tissues (Georgilis et al., 2018; Laberge et al., 2015; Lau and David, 2019; Lopes-Paciencia et al., 2019). As a remedy to senescence, we identify both classical and novel pathways associated with CEnC senescence that can be manipulated experimentally to potentially achieve significant CEnC expansion with minimal senescence. In addition, we identified which previously reported markers represent the optimal markers for selection of high quality cultured CEnC, and propose *TMEM178A* as a novel positive selection marker. We propose that because senescent CEnC can be present in cultured CEnC preparations and that senescent cells may support tumorigenesis, a Descemet membrane biomimetic carrier is preferred to injection of a suspension of cells, to both maximize cell count and minimize their tumorigenic potential (Gutermuth et al., 2019; Kinoshita et al., 2018; Teichmann et al., 2013). In conclusion, while additional changes to the dual media method may improve the culturing of CEnC in vitro, we provide evidence that this method is preferred to continuous culture in mitogen-rich media.

---

**Figure S1.**
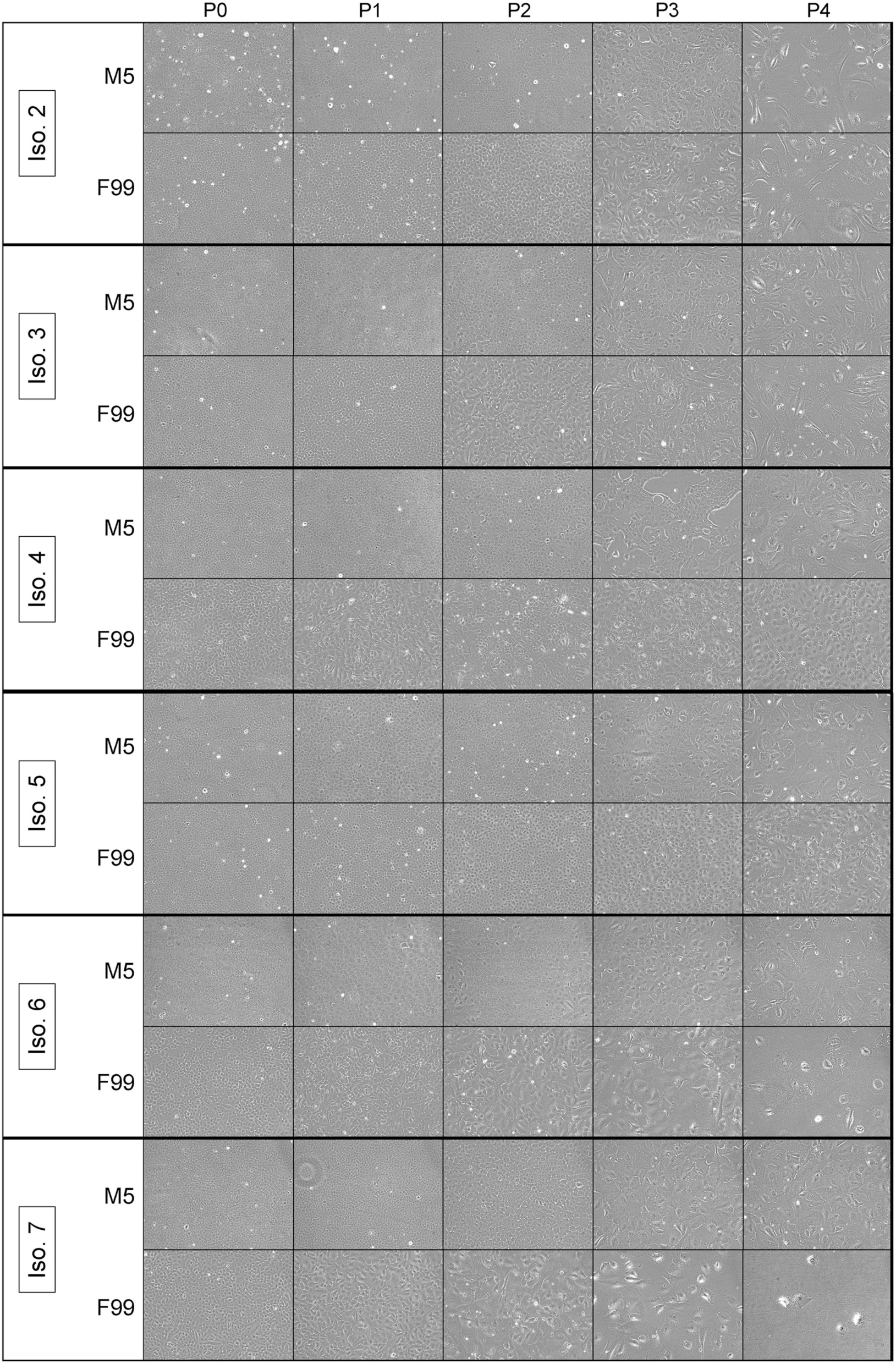
Phase contrast microscopy of additional independent CEnC cultures. Images show six additional independent CEnC cultures at 100% confluence for five passages (P0-P4). These six are in addition to the culture described in Figure 1.

**Figure S2.**
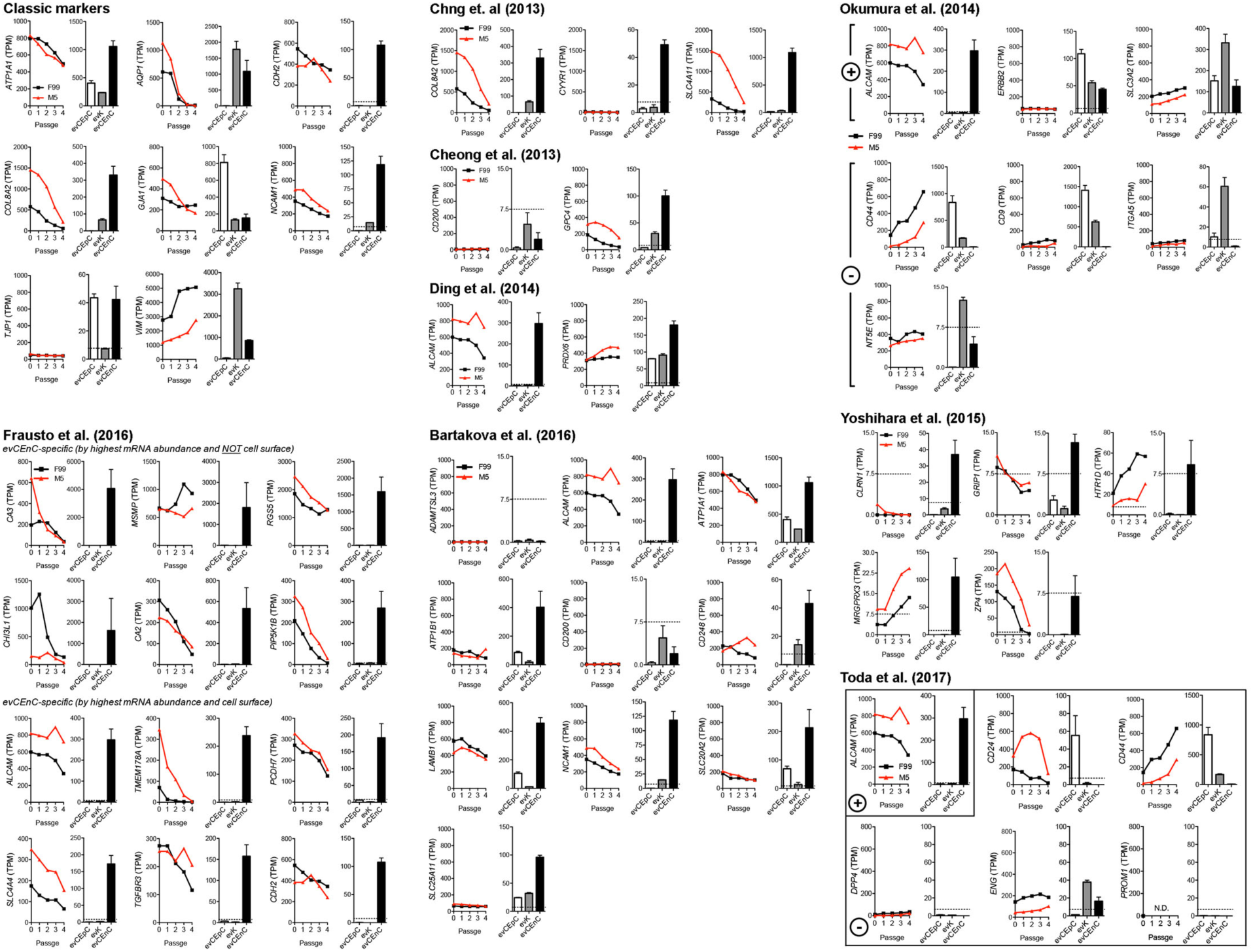
Published CEnC markers and the effect that passaging has on their expression. Analysis results for previously published CEnC markers using data from passaged CEnC and ex vivo data from the three main cell types of the cornea (epithelial cells (evCEnC), keratocytes (evK) and endothelial cells (evCEnC)). Plus and negative signs indicate studies that identified both positive and negative selection markers. Data in line graphs are represented as the mean TPM at each passage (n=7). Data in bar graphs are represented as the mean TPM±SEM (n=3). This analysis was performed for qualitative assessment of previously published markers.

**Figure S3.**
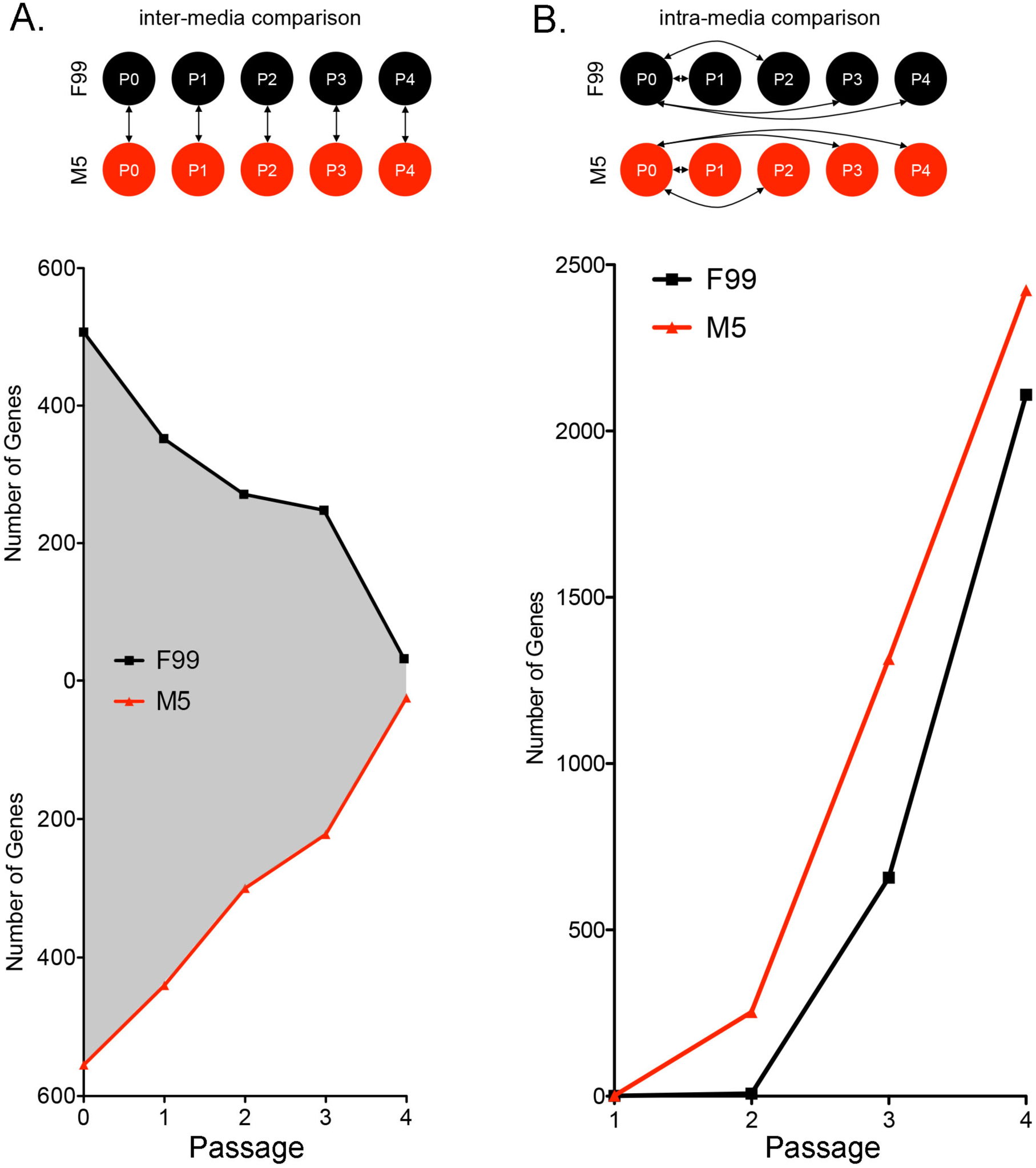
Number of identified differentially expressed genes from inter- and intra-media comparisons. (A) Schematic representing the inter-media comparisons (double arrows) made for the identification of differentially expressed genes (i.e., upregulated in F99 and upregulated in M5). Line graphs show the number of upregulated genes at each passage for both F99 and M5. The gray area between the curves represents the total number of upregulated genes, irrespective of media. At P0, approximately 1000 genes are upregulated in F99 and M5, and represents the largest difference for the F99 versus M5 comparisons. In addition, at P4, approximately 50 genes are upregulated in F99 and M5, and represents the smallest observed difference for the F99 versus M5 comparisons. (B) Schematic representing the intra-media comparisons (double arrows) made for the identification of differentially expressed genes at each passage (P0 was used as the reference for each comparison). Line graph showing the number of differentially expressed genes at each passage within each media (F99 and M5).

## SUPPLEMENTARY TABLES

**Table S1.** Donor cornea information

**Table S2.** Quantitative PCR primers.

**Table S3.** Antibodies.

**Table S4.** Differential gene expression lists for inter-media comparisons.

**Table S5.** Full results of gene ontology and pathway enrichment in inter-media data sets.

**Table S6.** Expression of cell senescence-associated genes in primary CEnC.

**Table S7.** Differential gene expression lists for intra-media comparisons.

**Table S8.** Full results of gene ontology and pathway enrichment in intra-media data sets.

**Table S9.** Full results of activation of Ingenuity canonical pathway in intra-media data sets.

**Table S10.** Full results of activation of Ingenuity cell cycle functions in intra-media data sets.

