## Supplemental Figures for "Phenotypic and functional characterization of corneal endothelial cells during in vitro expansion"

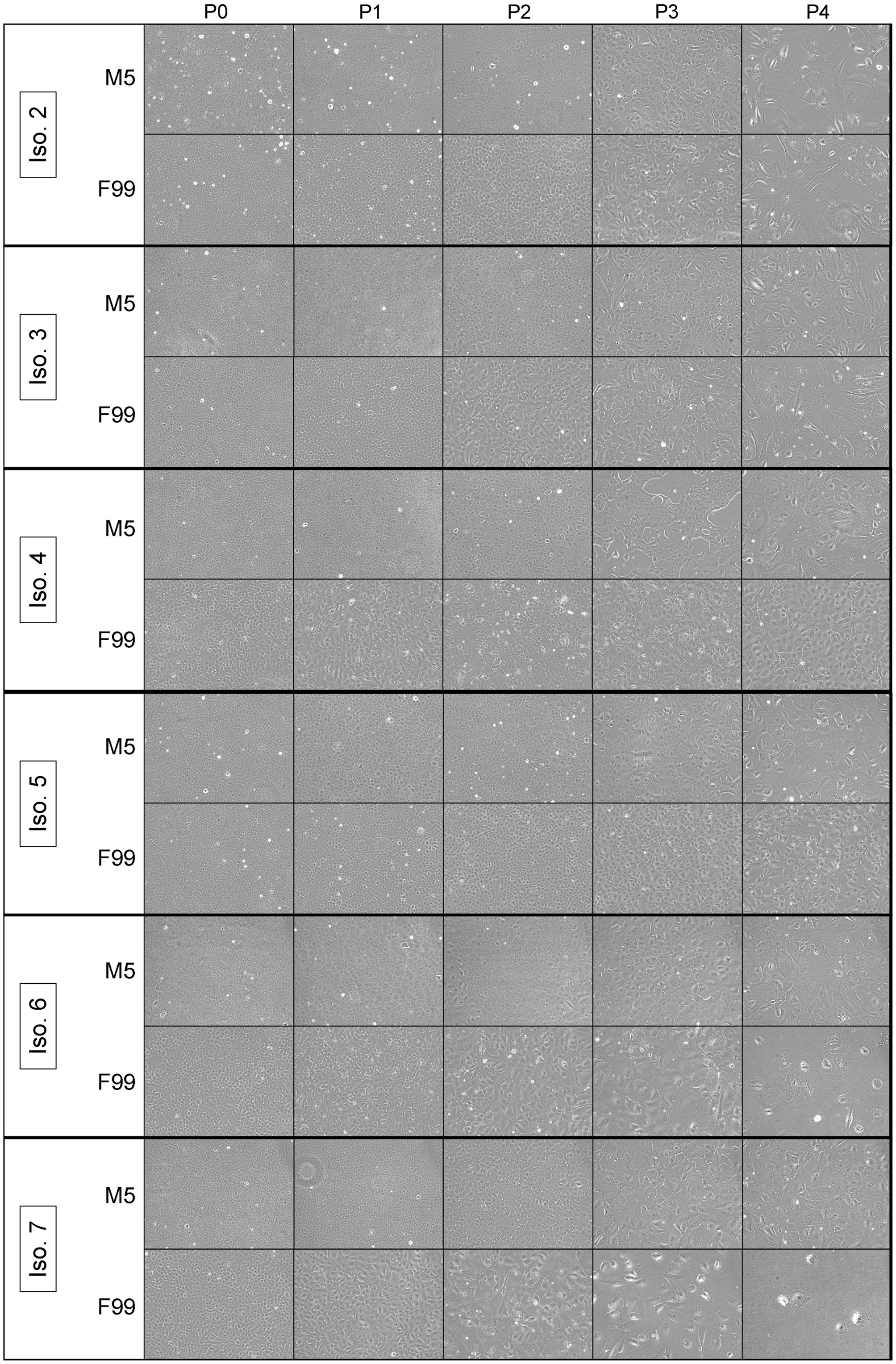


**Figure S1. Phase contrast microscopy of additional independent CEnC cultures.** Images show six additional independent CEnC cultures at 100% confluence for five passages (P0-P4). These six are in addition to the culture described in Figure 1.

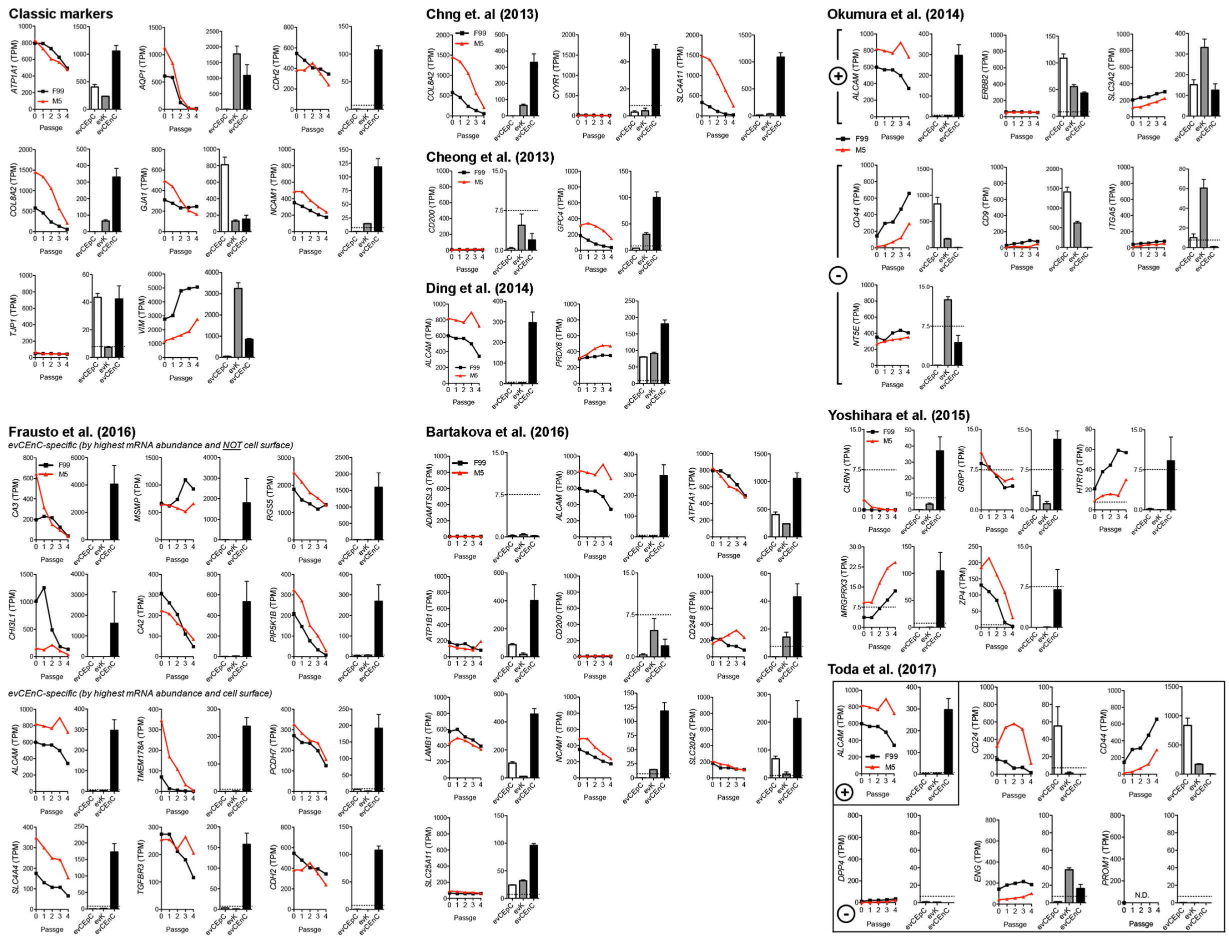


**Figure S2. Published CEnC markers and the effect that passaging has on their expression.** Analysis results for previously published CEnC markers using data from passaged CEnC and ex vivo data from the three main cell types of the cornea (epithelial cells (evCEnC), keratocytes (evK) and endothelial cells (evCEnC)). Plus and negative signs indicate studies that identified both positive and negative selection markers. Data in line graphs are represented as the mean TPM at each passage (n=7). Data in bar graphs are represented as the mean TPM±SEM (n=3). This analysis was performed for qualitative assessment of previously published markers.

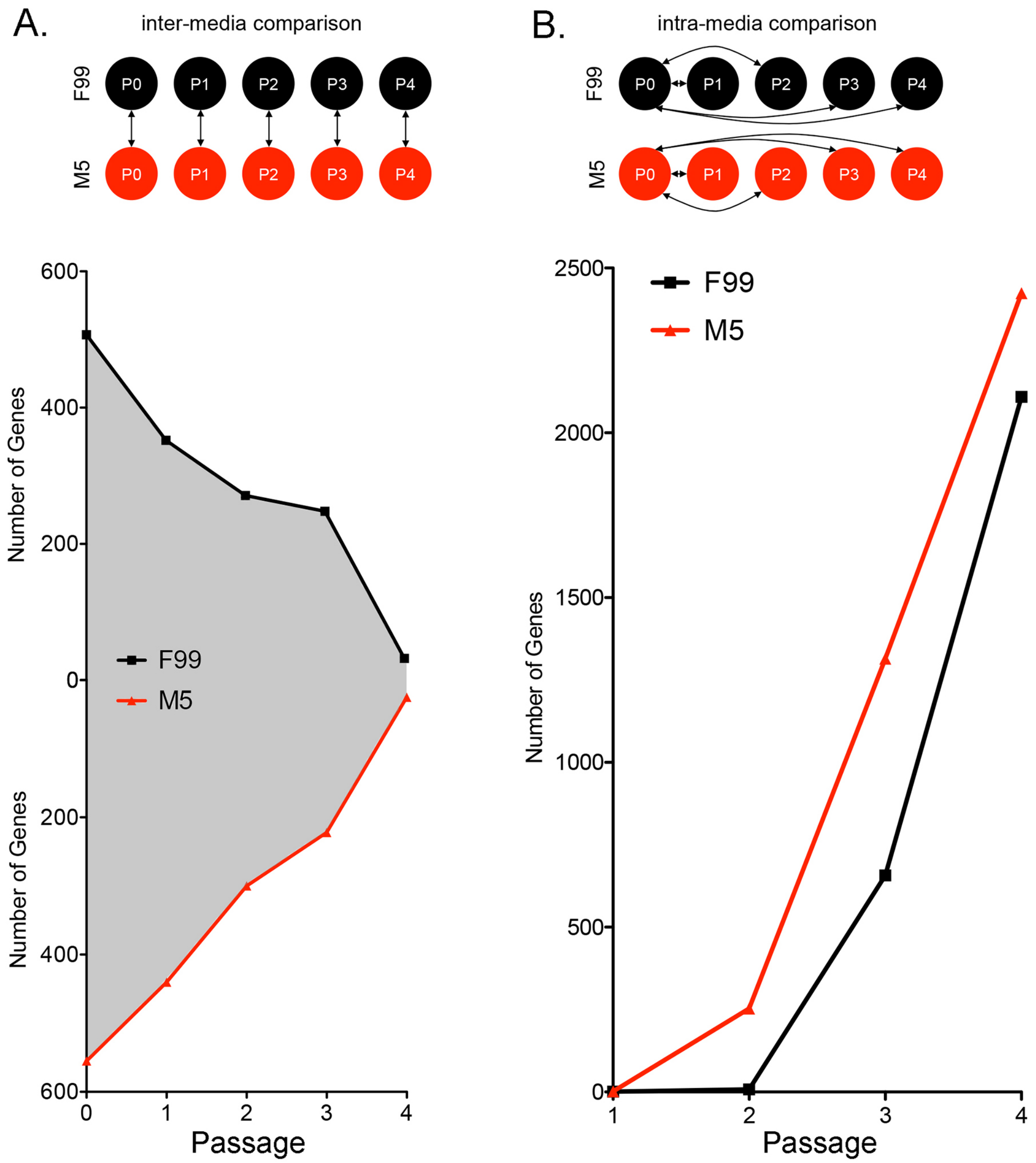


**Figure S3. Number of identified differentially expressed genes from inter- and intra-media comparisons.** (A) Schematic representing the inter-media comparisons (double arrows) made for the identification of differentially expressed genes (i.e., upregulated in F99 and upregulated in M5). Line graphs show the number of upregulated genes at each passage for both F99 and M5. The gray area between the curves represents the total number of upregulated genes, irrespective of media. At P0, approximately 1000 genes are upregulated in F99 and M5, and represents the largest difference for the F99 versus M5 comparisons. In addition, at P4, approximately 50 genes are upregulated in F99 and M5, and represents the smallest observed difference for the F99 versus M5 comparisons. (B) Schematic representing the intra-media comparisons (double arrows) made for the identification of differentially expressed genes at each passage (P0 was used as the reference for each comparison). Line graph showing the number of differentially expressed genes at each passage within each media (F99 and M5).
